# Gated feedforward inhibition in the frontal cortex releases goal-directed action

**DOI:** 10.1101/2020.12.13.422593

**Authors:** Jae-Hyun Kim, Dong-Hyun Ma, Eunji Jung, Ilsong Choi, Seung-Hee Lee

## Abstract

Cortical circuits process sensory information and generate motor signals in animals performing perceptual tasks. However, it is still unclear how sensory inputs generate motor signals in the cortex to initiate goal-directed action. Here, we identified a visual-to-motor inhibitory circuit in the anterior cingulate cortex (ACC) that induced action initiation in mice performing visual Go/No-go tasks. Interestingly, higher activity in sensory neurons and faster suppression in motor neurons of the ACC predicted faster reaction times. Notably, optogenetic activation of visual inputs in the ACC evoked strong suppression of neighboring motor neurons by activating fast-spiking sensory neurons and drove task-relevant actions in mice via activating striatal neurons. Finally, the ACC network activity maintained low during spontaneous and perceptual actions and increased during action cancellation in response to the stop signals. Collectively, our data demonstrate that visual salience in the frontal cortex exerts gated feedforward inhibition to release goal-directed actions.

## Introduction

The transformation of salient sensory stimuli into a goal-directed motor action is a fundamental process of the mammalian cortex^1^. In animals performing perceptual tasks, task-relevant activity has been identified in distinct cell types and circuits of the frontal cortex^2–4^. However, it is still unclear how these cell types and circuits causally interact to evoke goal-directed actions. In both rodent and primate frontal cortices, the ramp-to-threshold premotor activity has been shown to determine response times in animals making sensory-guided motor decisions^5,6^. Modeling studies have implied that stochastic accumulation of sensory evidence to the ramping activity is gated in order to generate timely goal-directed actions during perceptual decision-making^7,8^. However, empirical evidence of this gating model is lacking, and the neural circuit mechanisms of transferring discrete sensory information to ramping motor signals at different times in the frontal cortex remain unclear.

Among sub-regions in the frontal cortex, the anterior cingulate cortex (ACC) guides animals in different ways to perform goal-directed perceptual actions precisely. First, it prevents premature responses in the absence of the sensory cues^9^. Second, the ACC exerts the top-down modulation of the visual cortex (VC) to improve the perceptual process^10,11^. Finally, the ACC is involved in various functions that guide perceptual decisions by monitoring performance^12,13^, expecting rewards^14^, and controlling action selection^15^. In the mouse cortex, the VC projects to the ACC, which projects back to the VC, making a strong cortical loop^11,16^. This anatomical feature of the mouse ACC is similar to the primate frontal eye field (FEF), which also makes a strong reciprocal connection to the visual cortex^17^. The FEF not only exerts the top-down modulation of VC neurons^18^ but also encodes distinct sensory (visual) and motor (movement) signals when the animal performs a visually guided decision-making task^8,19^. In the moue ACC, however, it is unclear how distinct visual and motor signals appear in the ACC during the visuomotor transformation process. Although the frontal activity representing sensory and motor signals was examined extensively in animal models of perceptual decision-making, it is largely unknown how the bottom-up sensory inputs trigger motor signals in the frontal cortex to evoke relevant actions in task-performing mice.

In this study, we found that the ACC transforms visual information into decisive action in mice performing the visual Go/No-go (GNG) detection task: inactivating the ACC increased impulsive licking and disrupted the perceptual licking that happens after the visual cue. Using *in vivo* multichannel recordings, we identified two populations of neurons that show visual responses with gain modulation and ramp-to-threshold pre-licking activity in the ACC of task-performing mice. Interestingly, the amplitude in visual responses and the timing of pre-lick suppression significantly correlated with licking response times. Furthermore, the optogenetic activation of ACC sensory neurons that receive direct inputs from the VC was sufficient to suppress neighboring ACC neurons and generate licking action even in the absence of visual stimuli. Optogenetic activation of VC axon terminals in the ACC induced activation of response-increased neurons in the striatum. Therefore, the increase of sensory signals suppressed motor neurons in the ACC, and this sensory-to-motor suppression in turn released licking action via activating the basal ganglia loop. Interestingly, neighboring neurons, which were neither sensory nor motor, showed a tonic activity that inhibited sensory-to-motor suppression: when the neighboring activity was low, the sensory-to-motor suppression occurred. Our findings illustrate how neural circuits in the ACC transform salient visual cues into the release of motor actions. Importantly, ACC balances between the inhibition and the execution of goal-directed action by gating the sensory-induced suppression of the sustained motor activity that restrains impatient responses.

## Results

### ACC is involved in visually guided goal-directed behaviors

To examine how visual information is transformed into precise motor action, we trained head-fixed mice to perform a simple visual detection task using full-field flashing light stimuli (120 and 0 lux) under the GNG regime (Fig. 1a; Methods). When mice executed licking responses after the flashing visual stimuli in Go trials, we gave a water reward and counted the trial as ‘Hit’. Across the session, the licking action can happen either precisely as a controlled response to the visual stimuli or spontaneously as a premature response before the visual cue during the task. Novice mice constantly licked even before the introduction of visual stimuli in most hit trials (’impulsive lick’ hit trials (Hit_IL_); Fig. 1b). Trained mice, however, licked after the visual cues in most Go trials (’perceptual lick’ hit trials (Hit_PL_); Fig. 1c and Supplementary Video 1). During the training, constant gray stimuli (60 lux) were presented as background stimuli (BG), and the failure to withhold licking in the BG trial was counted as a false alarm (FA). Novice mice showed more FA than expert mice (Fig. 1b,c). Most mice became experts in performing the task within 10 sessions (Fig. 1d). To expert mice, we additionally presented weaker visual stimuli and measured their licking responses: Hit_PL_ rates gradually increased as a function of visual intensity, whereas Hit_IL_ rates remained low across the trials (Fig. 1e). These data indicate that the trained mice release their licks after visual perception and otherwise withhold licks.

**Fig. 1.**
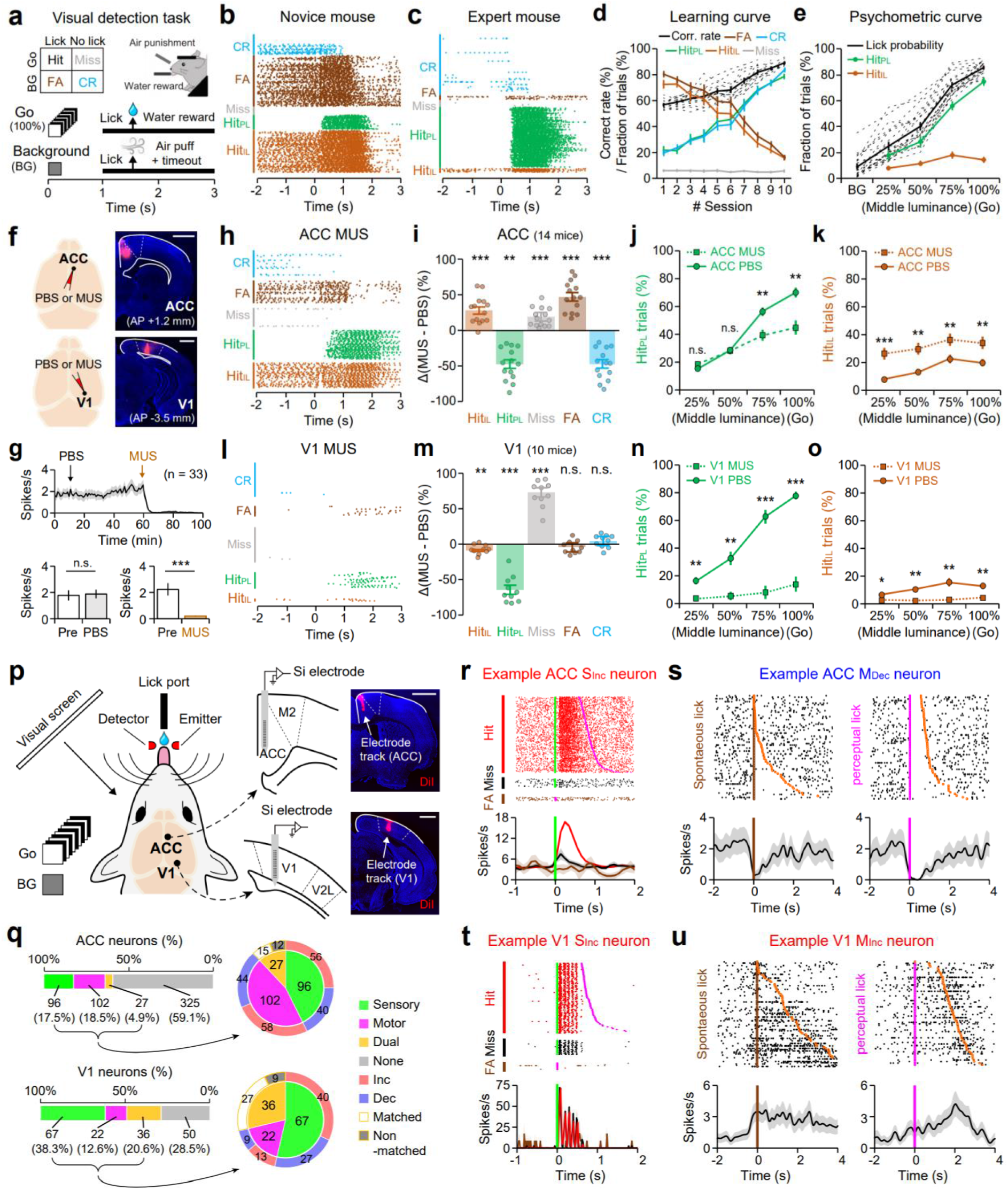
Functional roles of the ACC and the V1 that contain neurons representing sensory and motor signals in mice performing a visual Go/No-go task. **a**, Schematic illustration of the visual detection task. **b-e**, Task performance of trained mice across sessions. Colors represent trial types. Cyan, correct rejection (CR); brown, false alarm (FA); gray, Miss; green, Hit_PL_; light brown, Hit_IL_. **b**,**c**, Lick raster plots of a novice (**b**) and an expert mouse (**c**) during a session. Each dot represents a single lick. **d**, Learning curves that represent rates of each trial type across 10 consecutive sessions. Black lines, correct rates (dotted, individual mice; solid, average). **e**, Psychometric curves of expert mice. Licking rates across luminance levels of visual stimuli. Black lines, lick probabilities at each trial type (dotted, individual mice; solid, average). **f**, Schematic of PBS or MUS injection (left) and histological confirmation of MUS injection in the coronal sections of ACC and V1 (right). Scale bars, 1 mm. **g**, Effects of PBS and MUS injection in the ACC. Average firing rates (top), comparison of 10-min-averaged spike rates before (Pre) and after PBS (bottom left) or MUS (bottom right) injection. **h-k**, Effect of MUS injection in the ACC. **h**, Lick raster plot of an expert mouse injected with MUS. **i**, % changes of Hit_IL_, Hit_PL_, Miss, FA, and CR rates by MUS injection compared with PBS injection. **j**, % of Hit_PL_ across trials with different luminance stimuli. Dotted lines with squares, ACC MUS injection; solid lines with circles, ACC PBS injection. **k**, Same as **j**, but for Hit_IL_ trials. **l-o**, Same as **h-k**, but for MUS injection in the V1. **p**, Left, schematic of *in vivo* multichannel recordings in ACC or V1 of a task-performing mouse. Right, brain slices labeled with DiI on the electrode track in ACC (top) and V1 (bottom). Scale bars, 1 mm. **q**, Categorization of neuronal types in ACC (top; n = 550, 21 mice) and V1 (bottom; n = 175, 11 mice). Horizontal bars, all recorded neurons; pie charts, responsive neurons. Green, sensory; magenta, motor; yellow, dual-responsive neurons; gray, non-responsive neurons. Responsive neurons showed either increased (red) or decreased (blue) firing activity as a response. Dual neurons showed either the same (Matched) or the opposite (Non-matched) direction of activity changes between sensory and motor events. **r**, Visually-evoked activity of example ACC S_Inc_ (left) and S_Dec_ neurons (right). Raster plots (top) and PSTHs (bottom) from Hit (red for S_Inc_ and blue for S_Dec_), Miss (black), and FA trials (brown). Green and magenta ticks indicate stimuli and lick onset, respectively. **s**, Raster plots (top) and PETH (bottom, spikes/s) of an example ACC M_Dec_ neuron, aligned to the onsets of spontaneous licking (left, brown) and perceptual licking (middle, magenta). Orange, the offset of licking. **t**,**u**, Same as **r**,**s**, but for V1 S_Inc_ (**t**) and M_Iec_ neurons (u). n.s. (not significant), \**p* < 0.05, \*\**p* < 0.01, \*\*\**p* < 0.001; Wilcoxon signed-rank test with Bonferroni correction. Error bars show ± SEM.

To determine which cortical areas are involved in visual decision-making during the task, we pharmacologically silenced one of five sub-regions of the dorsal cortex on the contralateral hemisphere of an eye that had been presented with visual stimuli by injecting it with muscimol (MUS), a GABA_A_ receptor agonist (Fig. 1f and Supplementary Fig. 1a). MUS injection completely suppressed spiking activity in the injected area, whereas injection with phosphate-buffered saline (PBS) had no effect (Fig. 1g). Inactivation of the ACC, the primary visual cortex (V1), or the secondary visual cortex (medial area; V2M), but not the secondary motor cortex (M2) or the primary somatosensory cortex (S1), significantly impaired the performance of mice (Fig. 1h-o and Supplementary Fig. 1b-e). Surprisingly, inactivating ACC led to different consequences than inactivating V1 or V2M (Fig. 1i,m and Supplementary 1c_1_). ACC inactivation caused an increase in impulsive licking behavior and resulted in a significant increase of Hit_IL_ and FA trials as if expert mice became novice mice (Fig. 1h-k). On the other hand, silencing V1 or V2M caused a significant reduction of Hit_PL_ in all visual trials (Fig. 1n and Supplementary Fig. 1d_1_). Our data collectively indicate that the ACC, crucially, releases visually guided goal-directed actions in mice while withholding immature responses. In contrast, the VC is critical for detecting visual stimuli and initiating vision-guided action during the task.

### Sensory and motor-related signals appear in both the ACC and V1

We next measured the task-relevant neural activities by *in vivo* multi-channel recordings in cortical areas that we targeted with muscimol injections (Fig. 1p and Supplementary Fig. 7a). We identified sensory and motor neurons that showed significant activity changes in response to visual stimuli without licking and to licking without visual stimuli, respectively (Methods). Interestingly, the ACC represented sensory and motor signals equally well and independently in separate groups of neurons, having a small number of dual-responsive neurons (Fig. 1q, top). Compared with the ACC, the V1 and the V2M contained more sensory neurons, whereas the V1 and the M2 contained more dual-responsive neurons, having sensory responses that largely overlapped with motor-related activity (Fig. 1q (bottom) and Supplementary Fig. 7b). Furthermore, the activity of most of the dual-responsive neurons in the V1 and M2 showed corresponding increases or decreases during visual stimulation and licking (Fig. 1q and Supplementary Fig. 7b). In untrained mice, ACC visual responses were weak and contained a small number of sensory neurons, while their motor activity during licking was stronger than that of well-trained mice (Supplementary Fig. 2). This result suggests that sensory inputs to ACC are strengthened in a discrete neural population after task learning.

We next examined whether the visual response appears consistently across the trials regardless of licking action. Interestingly, ACC sensory neurons (both increased (S_Inc_) and decreased neurons (S_Dec_)) showed significantly enhanced visual responses during Hit_PL_ trials compared with miss trials (Fig. 1r and Fig. 2a-c). V1 neurons, however, showed similar visual responses between Hit_PL_ and miss trials, reliably relaying the physical attributes of visual stimuli (Fig. 1t and Supplementary Fig. 3a-c). We next found that ACC and V1 motor neurons showed significant changes in their activity (either increase (M_Inc_) or decrease (M_Dec_)) before the onset of spontaneous licking (−0.5 ∼ 0 s) and until the end of licking (Fig. 1s,u. Fig. 2d,e, and Supplementary Fig. 3d,e). Interestingly, ACC M_Dec_ neurons showed conserved premotor activity even before perceptual licking, whereas neither ACC M_Inc_ nor V1 M_Inc/Dec_ neurons did (Fig. 2f,g and Supplementary Fig. 3f,g). These data indicate that ACC M_Dec_ neurons represent the most reliable premotor signals of the lick decision in task-performing mice.

**Fig. 2.**
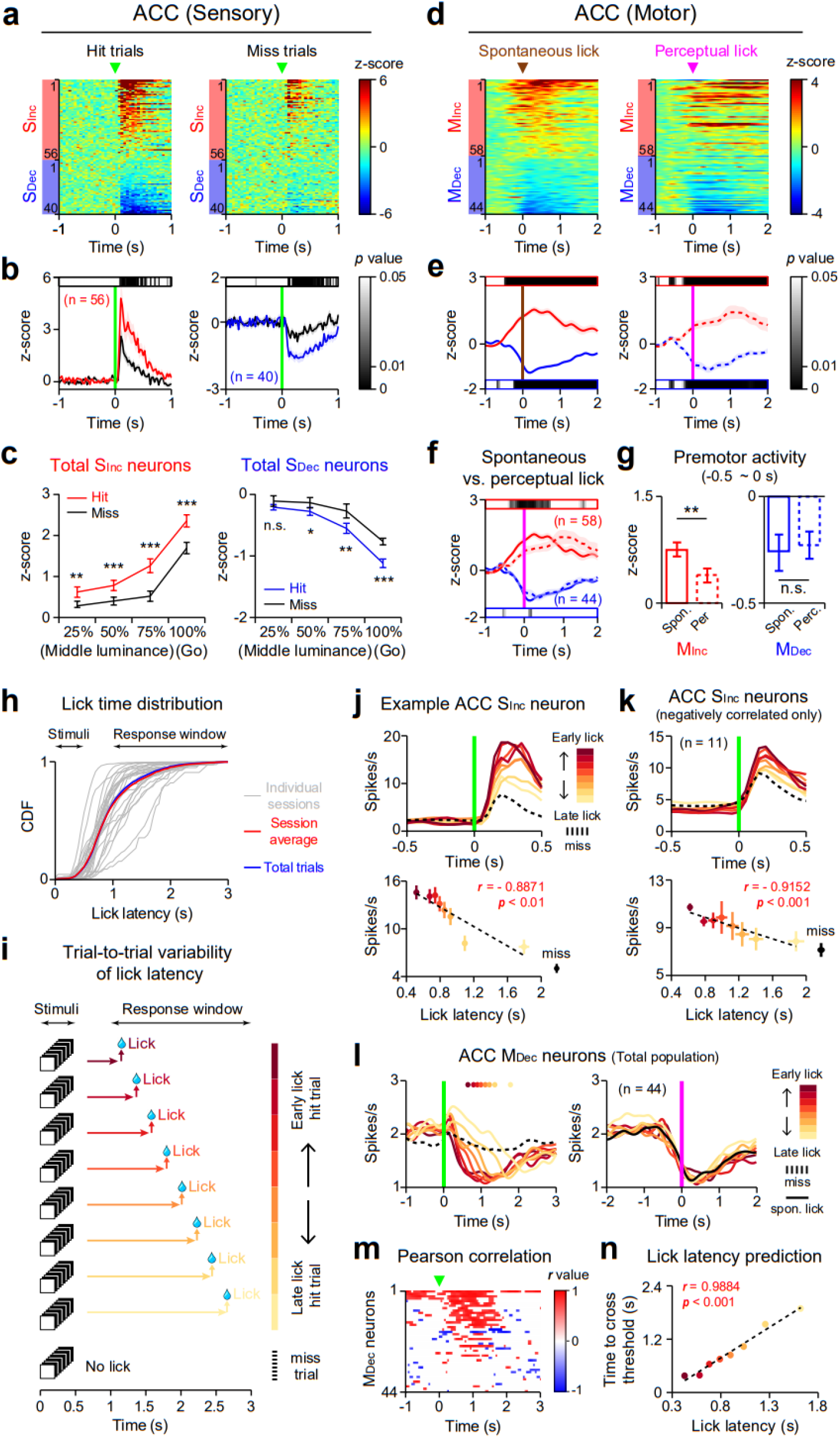
Response amplitudes of S_Inc_ neurons and times to reach a threshold in ramping-down activity of M_Dec_ neurons in the ACC predict trial-by-trial variations in response times of mice performing the task. **a**, Normalized and averaged population activity of ACC sensory neurons (n = 96) during hit (left) and miss trials (right). Neurons are sorted by hit trial responsiveness. **b**, Population average of normalized PSTHs from ACC S_Inc_ (left) and S_Dec_ neurons (right). Horizontal bars, color-coded *p*-values for significance tests at each time bin (25 ms). **c**, Population average of the mean visual signals (0 ∼ 0.5 s after stimuli onset) in ACC S_Inc_ (left) and S_Dec_ neurons (right) across different luminance levels of visual stimuli. **d**,**e**, Population activity (**d**) and normalized PETHs or PSTHs (**e**) of ACC M_Inc_ (red, n = 58) and M_Dec_ neurons (blue, n = 44) aligned to the onsets of spontaneous licking (left, brown), perceptual licking (middle, magenta), and stimuli onset (right, green). Horizontal bars above (red) and below (blue) PETHs or PSTHs denote color-coded *p*-values from the significance test with zero in ACC M_Inc_ and M_Dec_ neurons, respectively. **f**, Averaged PETHs of ACC M_Inc_ (red) and M_Dec_ neurons (blue) during spontaneous (solid lines) and perceptual licking (dotted lines). Horizontal bars denote color-coded *p*-values for significant differences in the activity of ACC M_Inc_ and M_Dec_ neurons at each time bin (50 ms) between spontaneous and perceptual licking. **g**, Premotor activities (−0.5 ∼ 0 s from licking onset) in ACC M_Inc_ (left) and M_Dec_ neurons (right) during spontaneous and perceptual licking. **h**, Distribution of lick response times in each session (gray), average across sessions (n = 26, red), and in total trials (n = 6265, blue). **i**, Schematic illustration of the trial-to-trial variation in lick latencies across hit trials. **j**, Top, color-coded PSTHs of an example ACC S_Inc_ neuron. Firing rates were averaged in trials sorted according to lick latencies (darker color, earlier lick trials). Dotted line, PSTH of the same neuron in miss trials. Bottom, scatter plots of lick latencies (x-axis) and visually evoked firing rates (y-axis) averaged across trials. Note the significant linear relationship between them. *r*, Pearson correlation coefficient; *p*, the significance of the correlation; dotted line, linear regression. **k**, Same as **j**, but for ACC S_Inc_ neurons with a significant negative correlation of their activity with lick latencies (neurons in the red box of Supplementary Fig. 5a). **l**, Color-coded PSTHs aligned to the onset of visual stimuli (left) and PETHs aligned to the onset of licking (right) of ACC M_Dec_ neurons in trials sorted as shown in **j**. Color-coded dots indicate average lick latencies from each segment of lick latency. **m**, The Pearson correlation coefficient of spiking activity aligned to the onset of stimuli (green arrowheads) and trial-by-trial lick latencies in ACC M_Dec_ neurons. Only significant bins were colored based on *r* values. Note a strong positive correlation between post-stimulus activities in M_Dec_ neurons and lick latencies. **n**, Pearson correlation (*r*) between lick latencies and times to cross lower thresholds in the activity of ACC M_Dec_ neurons. Error bars show ± SEM.

Since V1 premotor activity was significantly reduced during perceptual licking, we examined whether this activity is correlated with other orofacial movements, not just licking in head-fixed mice (Supplementary Fig. 4a-c and Supplementary Video 1). We found that pupil size, nose movement, and whisker movement increased together with spontaneous licking (Supplementary Fig. 4d). However, during perceptual licking, pupil size decreased while nose movement and whisking still occurred (Supplementary Fig. 4e). As V1 premotor activity was reduced significantly during perceptual licking (Supplementary Fig. 3f,g), it is highly possible that V1 premotor signals may be related to the pupil dilation of the contralateral eye.

### Visual amplitudes of S_Inc_ neurons and ramp-to-threshold premotor suppression of M_Dec_ neurons in the ACC predict response times

During the task, individual mice showed trial-to-trial variations in lick latency across hit trials (Fig. 2h). We examined whether the sensory and motor signals in the ACC can predict response times of perceptual licking (Fig. 2i). We found that ACC S_Inc_ neurons, but not S_Dec_ neurons, showed a significant correlation between the amplitude of visual responses and lick latency (Fig. 2j,k and Supplementary Fig. 5a,c,d). The larger the visual response of the ACC S_Inc_ neurons, the faster perceptual licking happened (Fig. 3j,k and Supplementary Fig. 5c). This gain modulation of the sensory signal was not observed in the V1, V2M, or M2 (Supplementary Fig. 5e-g and Supplementary Fig. 7c-e). Thus, the visual gain in ACC S_Inc_ neurons correlates with the speed of visual-to-motor transformation during the task and predicts response times.

**Fig. 3.**
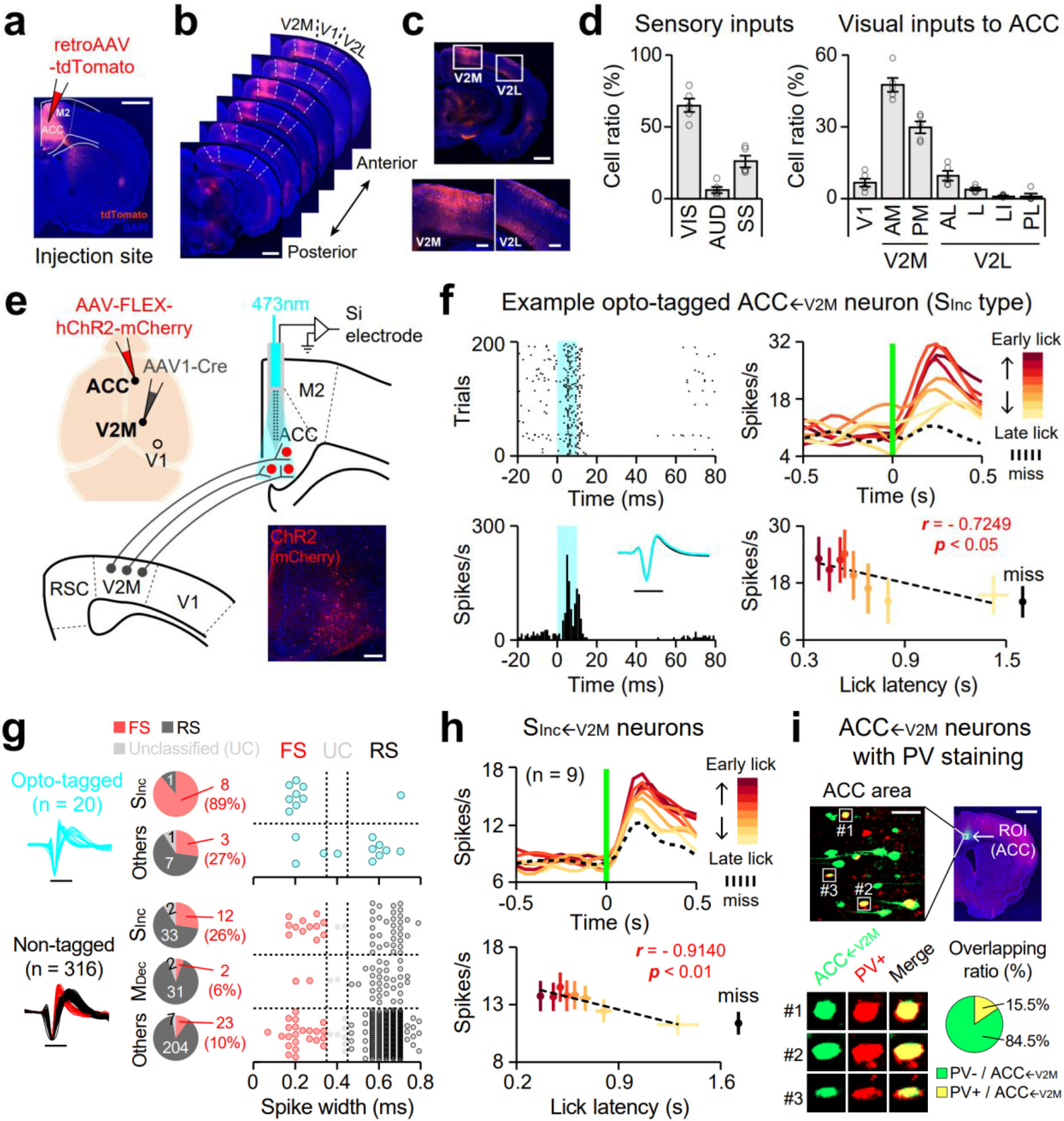
Optogenetic identification of ACC_←V2M_ neurons representing fast-spiking S_Inc_ neurons. **a**, Schematic diagram illustrating retrograde tracing of input neurons to the ACC. Scale bar, 1 mm. **b**, Coronal brain slices along anterior to posterior axis covering visual cortical areas. Scale bar, 1 mm. **c**, A coronal brain section including the V2M and V2L (top), and magnified images of the V2M (bottom left) and V2L (bottom right). Scale bars, 1 mm (top) and 200 μm (bottom). **d**, The ratio of labeled neurons quantified in the visual (VIS), auditory (AUD), and somatosensory (SS) cortices (left) and across sub-regions of the visual cortex (right). Abbreviations of the sub-regions are listed in Supplementary Table 2. Bars, mean ± SEM; total 5 mice. **e**, Schematic illustration of the opto-tagging identification of ACC_←V2M_ neurons. Bottom-right, histological confirmation of the ChR2-mCherry expression in the ACC. Scale bar, 200 μm. **f**, Left, spike raster (top) and PSTH (bottom) of a representative opto-tagged ACC_←V2M_ neuron. Note the neuron is directly activated by blue-light photostimulation (cyan, 10 ms). Inset, averaged spike waveforms of the neuron measured during spontaneous firing (black) or during optical stimulation (cyan). Scale bar, 1 ms. Right, color-coded PSTHs of an example opto-tagged ACC_←V2M_ neuron (S_Inc_ type) across trials sorted by lick latencies. **g**, Combined analysis of spike waveforms and task-relevant responses of opto-tagged and non-tagged neurons. Pie charts indicate fraction of neurons with distinct spike waveforms (red, fast-spiking (FS); dark gray, regular-spiking (RS); light gray, unclassified (UC); cyan, opto-tagged). Neuronal types were classified according to their task-relevant responses (S_Inc_, M_Dec_, and Others) and optically driven responses (top, opto-tagged; bottom, non-tagged). Dots, spike widths of individual neurons. **h**, Color-coded PSTHs of opto-tagged S_Inc←V2M_ neuron across trials sorted by lick latencies. Note the significant correlation between visually evoked activities and lick latencies. Error bars, ± SEM. **i**, Identification of PV+ neurons among ACC_←V2M_ neurons. Top, a coronal brain slice of the ACC. V2M recipient neurons (green) were immunostained for PV (red). Scale bars, 50 μm (left) and 1mm (right). Bottom left, three representative views of PV+ ACC_←V2M_ neurons. Bottom right, a fraction of PV+ (yellow) and PV-negative (green) ACC_←V2M_ neurons.

We next analyzed premotor signals in ACC motor neurons. Surprisingly, ACC M_Dec_ neurons, but not M_Inc_ neurons, displayed ramping-down activity up to a constant threshold at a constant rate before the licking was released (Fig. 2l and Supplementary Fig. 6a-e). About half of ACC M_Dec_ neurons showed a significant correlation between lick latency and times when the activity of motor neurons crossed the thresholds set at various levels (Supplementary Fig. 6a). Analyzing the Pearson correlation, we found that a substantial number of ACC M_Dec_ neurons, but not M_Inc_ neurons, showed a strong and significant correlation between lick latency and neural activity at each time point (Fig. 2m and Supplementary Fig. 6f). Unlike in the ACC, premotor suppression in the V1 did not show any correlation with lick latency (Supplementary Fig. 6a,h-m). Rather, the M_Inc_ neurons in the M2 showed a ramping-up premotor activity, which significantly correlated with lick latency (Supplementary Fig. 7f,g). Taken together, our results indicate that the ACC shows a unique feature of premotor signals that are represented by M_Dec_ neurons, which predict lick response times by showing ramping-down-to-threshold activity before licking.

### Fast-spiking neurons in the ACC receive direct synaptic inputs from the VC

To reveal visual inputs to the ACC sensory neurons, we traced neurons that project to the ACC by using the retrograde AAV expressing tdTomato (Fig. 3a). Consistent with recent studies^16^, VC neurons, particularly the V2M neurons, project onto the ACC more than other sensory cortical areas (Fig. 3b-d; abbreviation in Supplementary Table 2). We labeled postsynaptic neurons of the V2M in the ACC (ACC_←V2M_) neurons with Cre by injecting the AAV1-hSyn-Cre into the V2M^20^ and expressed those neurons with the channelrhodopsin-2 (ChR2) by additional injection of the AAV2-DIO-channelrhodopsin-2 (ChR2) into the ACC (Fig. 3e). Using the silicon electrode integrated with an optic fiber, we identified the ChR2-tagged ACC_←V2M_ neurons based on their light-driven responses (Fig. 3e,f; Methods). Interestingly, 45% (n = 9) of total 20 opto-tagged ACC_←V2M_ neurons were S_Inc_ neurons (Fig. 3g, top), whereas only 14.9% (n = 47) of total 316 non-tagged ACC neurons were S_Inc_ type (Fig. 3g, bottom). The opto-tagged S_Inc_ neurons (S_Inc←V2M_) displayed a significant linear relationship between the visual amplitude and lick latency, and none of the opto-tagged ACC_←V2M_ neurons were motor neurons (Fig. 3g,h). These data indicate that the visual inputs from the VC mainly innervate and drive S_Inc_ neurons in the ACC.

We further found that 55% of the opto-tagged ACC_←V2M_ neurons were fast-spiking (FS) with narrow waveforms (Fig. 3g, top). In particular, opto-tagged S_Inc←V2M_ neurons were predominantly FS (89%), whereas only 11.7% of non-tagged ACC neurons were FS (Fig. 3g, bottom). Like S_Inc←V2M_ neurons, all S_Inc_ neurons identified by blind recordings in task-performing mice contained more FS neurons than other types (Fig. 6a,d,g,j). Thus, FS neurons were the major recipient of the visual inputs, which represent S_Inc_ signals in the ACC. Likewise, when we immunostained ACC_←V2M_ neurons with antibodies against the parvalbumin (PV), a well-known calcium-binding protein that is mainly expressed in FS neurons, we found that 15.5%, more than the known fraction in the cortex (∼7-8%), were PV-positive (Fig. 3i)^21,22^.

### Optogenetic activation of ACC_←V2M_ neurons induces licking behavior by suppressing neighboring ACC neurons

We next wondered whether the FS S_Inc_ neurons in the ACC exert feedforward inhibition on the M_Dec_ neurons and release the visually induced licking in task-performing mice. We optogenetically activated either the soma of ACC_←V2M_ neurons or the axon of V2M neurons in the ACC at the same frequency as actual visual stimuli in random trials during the task (Fig. 4a). Interestingly, the photoactivation of V2M-recipient neurons or V2M axon terminals in the ACC was sufficient to induce licking, whereas light stimulation without ChR2 expression showed no effect (Fig. 4b-d). Activation of V2M axons caused weaker suppression of ACC neurons than the soma activation, and in correlation with this, photoactivation of soma induced faster licking than that of axons in the ACC (Fig. 4e). These data indicate that visual inputs to the ACC initiate licking action during the visual detection task.

**Fig. 4.**
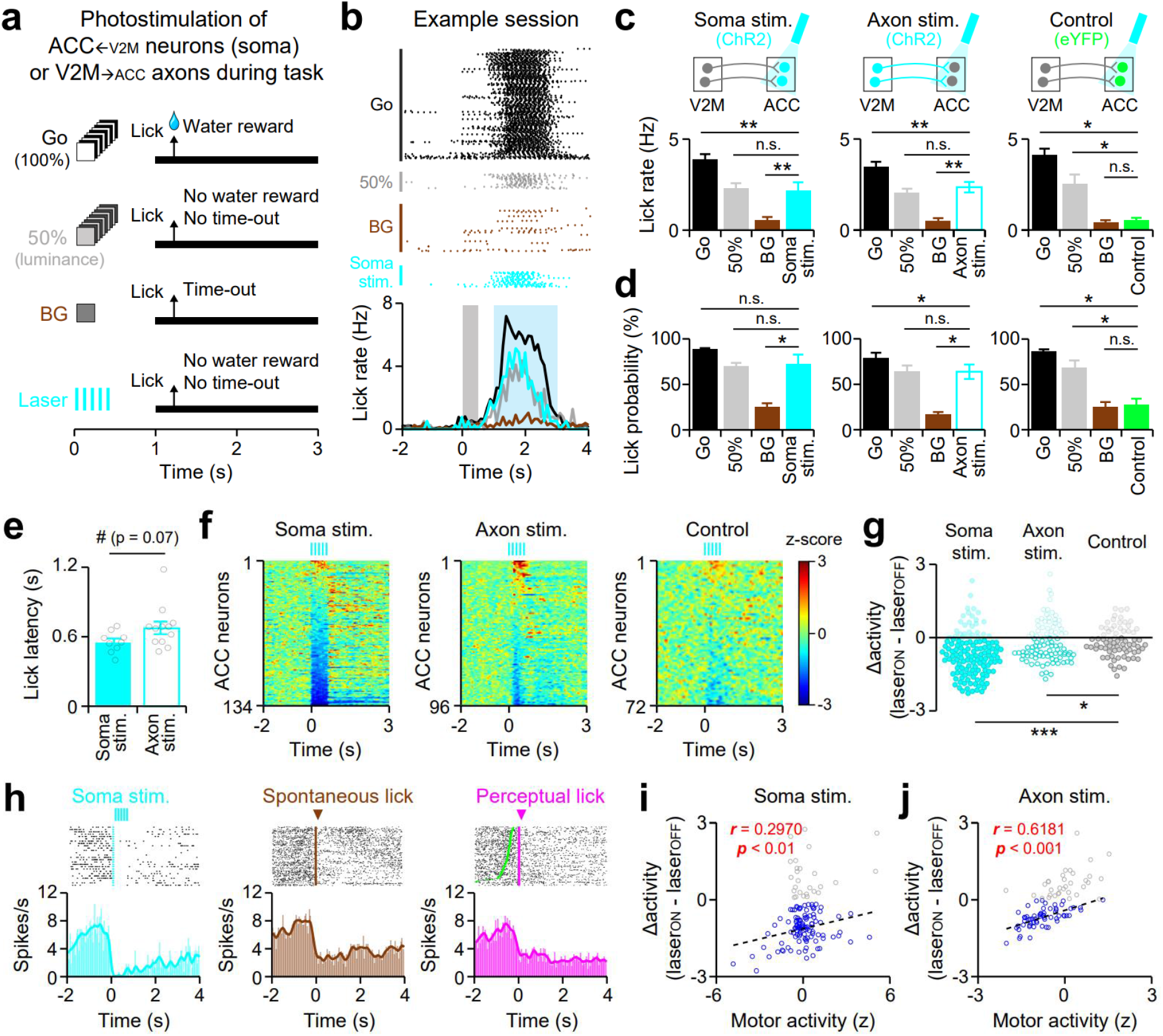
Optogenetic activation of V2M inputs to the ACC drives licking responses. **a**, Schematic of the visual detection task with different types of trials including photostimulation of the soma of ACC_←V2M_ neurons or V2M axon terminals in the ACC. Note that the frequency of the visual stimuli was identical to that of the blue-laser photostimulation. **b**, Lick raster (top) and lick rate histogram (bottom) of a mouse across trial types in an example session. Black, Go trials (100% luminance); gray, trials with visual stimuli with 50% luminance; brown, BG trials without visual stimuli; cyan, trials with photostimulation; gray shade, visual or photostimulation; blue shade, lick response window. **c**,**d**, Lick rate (**c**) and probability (**d**) from photostimulation of the soma of ACC_←V2M_ neurons expressing ChR2 (left, soma stimulation, 9 sessions using 3 mice), the axon of V2M neurons expressing ChR2 (middle, axon stimulation, 12 sessions using 3 mice), and the control group expressing eYFP in the soma of ACC_←V2M_ neurons (right, eYFP, 7 sessions using 3 mice). **e**, Lick latency from the onset of photostimulation. Gray open circles indicate data from individual sessions. **f**, Activity changes in ACC neurons by photostimulation of the soma of ACC_←V2M_ neurons (left), the axon of V2M neurons (middle) and the control group (right). **g**, Normalized activity (z) changes (Δactivity, laser_ON_ – laser_OFF_) by photostimulation of the soma of ACC_←V2M_ neurons (left, filled cyan circles), the axon of V2M neurons (middle, open cyan circles), and the control mice (right, gray circles). Statistical comparison was performed only among the suppressed neurons (negative Δactivity, dark circles). **h**, Spike raster (top) and mean firing rates (bottom) of an example ACC M_Dec_ neuron aligned to the onsets of photostimulation (left), spontaneous licking (middle), and perceptual licking (right). Note that the photostimulation of ACC_←V2M_ neurons suppressed M_Dec_ neuron. **i**, Scatter plot showing motor-related activity (z) and activity changes with the photostimulation of the soma of ACC_←V2M_ neurons (Δactivity, Laser_ON_ – Laser_OFF_). Pearson correlation (*r*) between the motor-related activity and the laser-evoked activity was calculated only in ACC neurons that were suppressed by the photostimulation (blue, n = 62). *r*, Pearson correlation coefficient; *p*, the significance of correlation; dotted lines, linear regression. **j**, Same as **i**, but for axon stimulation. n.s. (not significant), #*p* < 0.10, \**p* < 0.05, \*\**p* < 0.01, \*\*\**p* < 0.001; Wilcoxon sign-rank test (**c**, **d**); Wilcoxon rank-sum test (**e**, **g**). Error bars, ± SEM.

Not only did it induce licking, photostimulation of V2M inputs in the ACC suppressed the neural activity of neighboring neurons, including M_Dec_ neurons (Fig. 4f-h). Interestingly, the level of suppression by photostimulation in ACC neurons was significantly correlated with the level of reduction in their firing rates during licking (Fig. 4i,j). Taken together, our results demonstrate that the flow of visual information from V2M to ACC, triggered by the visual cues, evokes the motor response (licking) via feedforward inhibition; this process suppresses the activity of M_Dec_ neurons to a threshold at which the licking action is released.

### Optogenetic activation of V2M inputs in the ACC activates neurons in the striatum

Previous studies found that the ACC strongly projects to the dorsomedial striatum (DMS)^23^, and the ACC-to-DMS circuit might be important for inhibitory control of an action^24^. We also found a robust projection from the ACC to the DMS with retrograde tracing (Fig. 5a,b). Interestingly, these DMS-projecting ACC neurons rarely overlapped with the ACC_←V2M_ neurons (Fig. 5a,b), suggesting that the DMS-projecting ACC neurons are more related to the motor than to the sensory neurons.

**Fig. 5.**
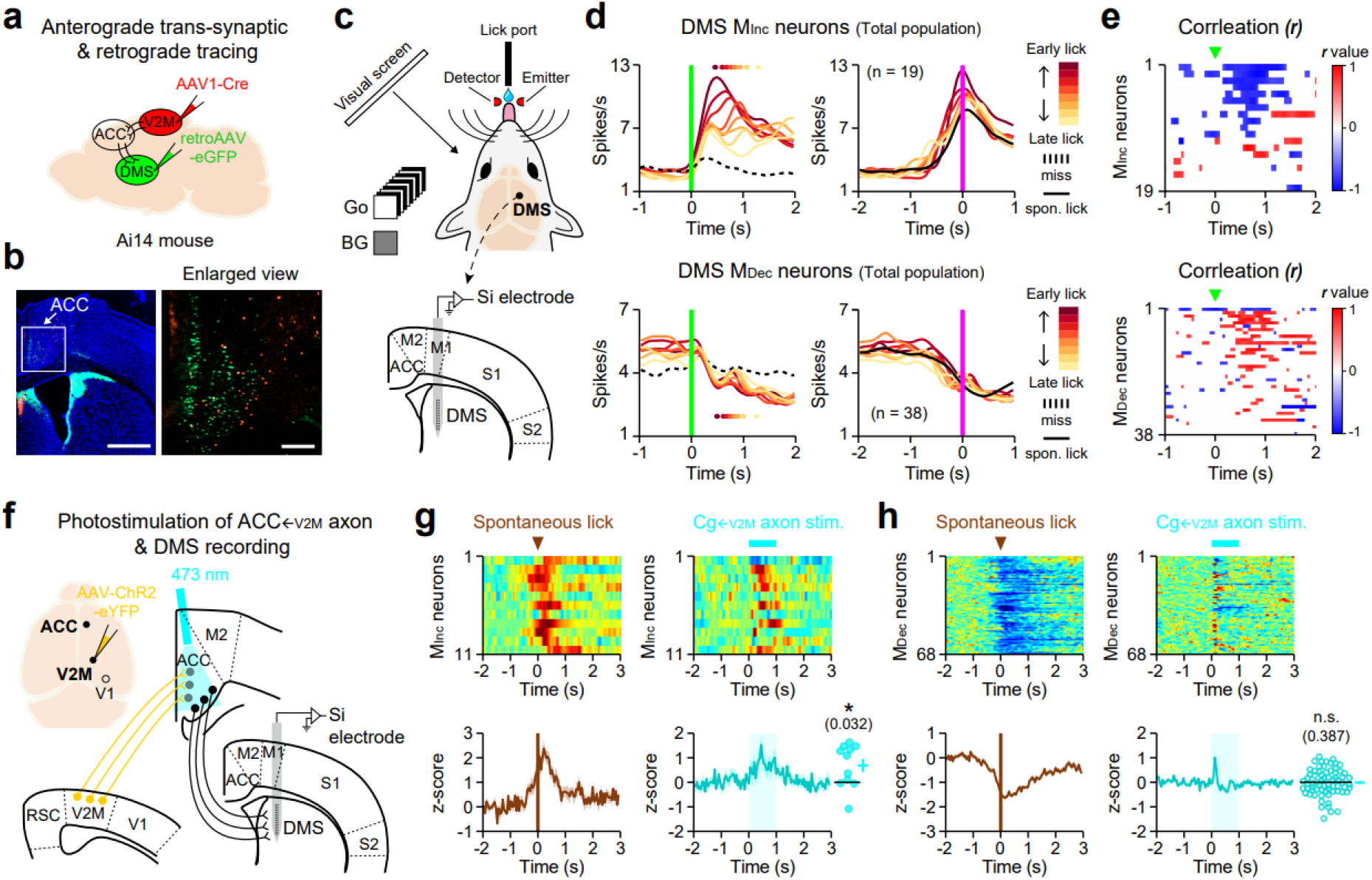
Photostimulation of V2M axons in the ACC activates task-relevant motor neurons in the DMS. **a**. Schematic of the DMS retrograde tracing with V2M anterograde trans-synaptic tracing. **b**. DMS-projecting (green) and V2M-recipient cells (red) in a wide view of the ACC area (left) and enlarged view of ACC area (right, white box in the left image). Scale bars, 1 mm (left) and 200 μm (right). **c**, Schematic of DMS activity recording during the visual detection task. **d**,**e**, Same as Fig. 2**l,m** but for DMS M_Inc_ (top, n = 19) and M_Dec_ neurons (bottom, n = 38). **f**, Schematic of DMS activity recording with photostimulation of V2M axons in the ACC. **g**, z-scored population activity heat-map (top) and average trace (bottom) of DMS M_Inc_ neurons (n = 11) during spontaneous lick (left) and photostimulation of ACC_←V2M_ axons. Bottom right inset, neural activity change by photostimulation of ACC_←V2M_ axons. **h**, Same as **f**, but for DMS M_Dec_ neurons (n = 68). n.s. (not significant), \**p* < 0.05; Wilcoxon sign-rank test. Error bars, ± SEM.

We next measured neural activity in the DMS, the downstream motor region of the ACC, during the visual detection task (Fig. 5c). We identified DMS neurons that showed a significant increase (M_Inc_ neurons, n = 11) or decrease (M_Dec_ neurons, n = 68) in their firing activity before licking. Interestingly, both the time required to cross the threshold and the amplitude of the activity of M_Inc_ but not M_Dec_ neurons in the DMS showed a significant correlation with the lick latency in task-performing mice (Fig. 5d,e). We further found that the optical activation of V2M axons in the ACC, which induced ACC M_Dec_ signal (Fig. 4f,g,j), also induced lick-related M_Inc_ signal in the DMS (Fig. 5g-h). Activity in DMS M_Inc_ neurons may initiate licking action via a direct pathway in the basal ganglia. Together, our data support the idea that suppression of M_Dec_ neurons in the ACC is a key process of sensory-to-motor transformation, releasing licking responses via activating downstream motor areas such as the basal ganglia during the visual GNG task.

### Network activity in the ACC gates sensory-to-motor suppression to release spontaneous licking

To understand how premotor suppression occurs consistently in M_Dec_ neurons, even without the sensory drive during licking without visual stimuli (spontaneous licking), we examined the activity of S_Inc_ neurons in the ACC during spontaneous licking. Interestingly, S_Inc_ neurons as well as opto-tagged S_Inc←V2M_ neurons showed a small but significant increase in their activity before the suppression of M_Dec_ neurons and spontaneous licking (Fig. 6a-f), suggesting that the process of feedforward inhibition was triggered by S_Inc_ neurons during spontaneous licking. However, during miss trials, although the visually induced activity of S_Inc_ neurons exceeded their activity during spontaneous licking, the suppression of M_Dec_ neurons did not happen (Fig. 6b). Thus, the simple feedforward inhibition does not fully explain the process of M_Dec_ suppression by FS S_Inc_ neurons (sensory-to-motor suppression); the gating process may exist to permit a small increase in S_Inc_ neurons to suppress M_Dec_ neurons during spontaneous licking but not during miss conditions.

**Fig. 6.**
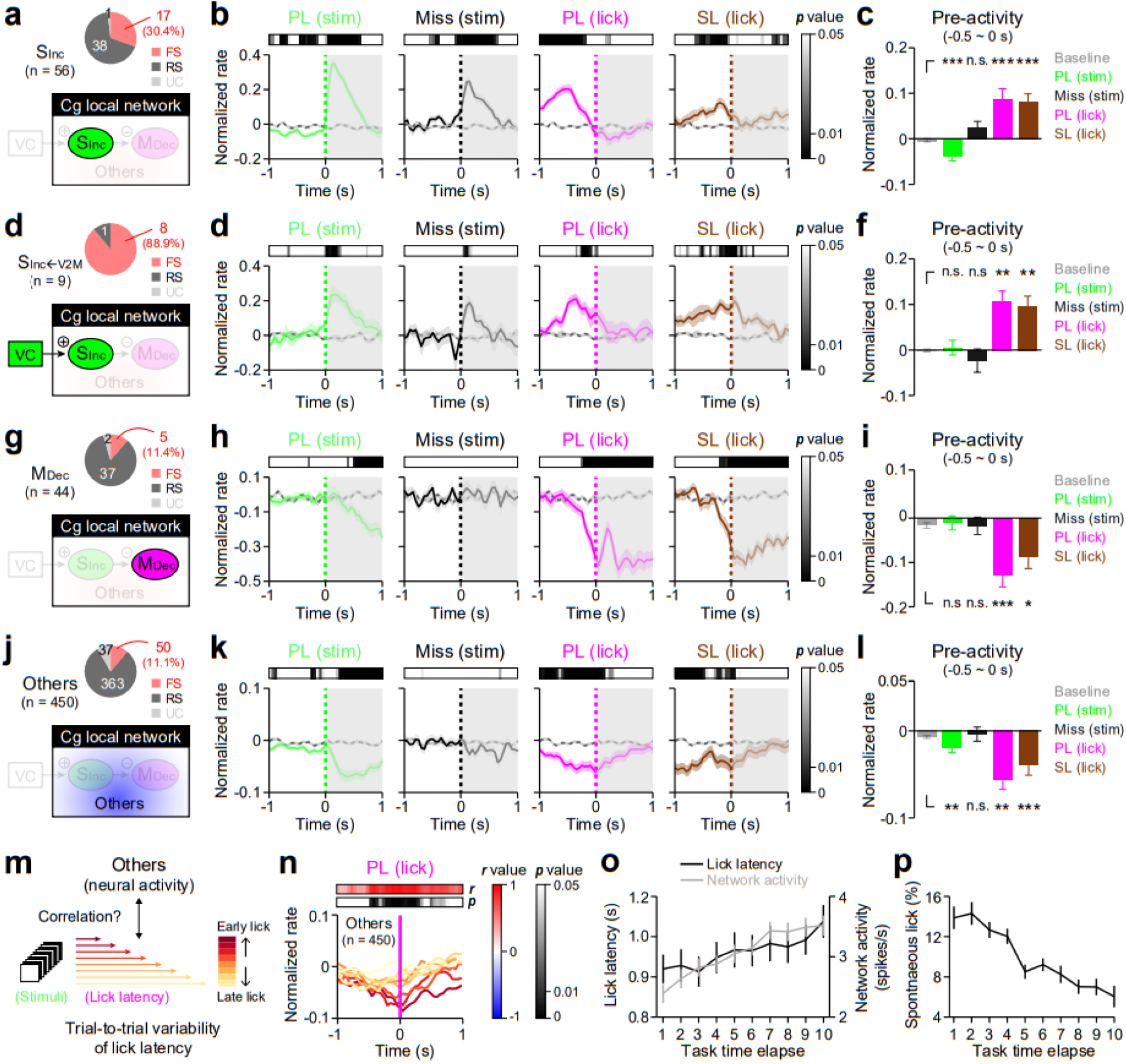
Neighboring neurons in the ACC gate sensory-to-motor suppression. **a**, Fraction of FS, RS, and UC neurons among ACC S_Inc_ neurons. **b**, Normalized firing rates of ACC S_Inc_ neurons aligned to the onset of stimuli during perceptual licking (green, PL (stim)) and during miss trials (black, Miss (stim)), aligned to the onset of licking during perceptual licking (magenta, PL (lick)) and during spontaneous licking (brown, SL (lick)). Firing rates were normalized by baseline activity (gray). Horizontal bars, color-coded *p*-values showing significant difference from baseline activity. **c**, Mean pre-activity of ACC S_Inc_ neurons (−0.5 ∼ 0 s from the onset of events). **d-l**, Same as **a-c**, but for opto-tagged S_Inc←V2M_ neurons (**d-f**), ACC M_Dec_ neurons (**g-i**), and other ACC neurons that are neither S_Inc_ nor M_Dec_ neurons (**j-l**). **m**, Schematic illustration showing the correlation between neural activities of neighboring neurons in the ACC (Others) and lick latencies. **n**, The Pearson correlation coefficient (*r*) and significance (*p*) between lick latencies and the normalized firing rates of ACC neighboring neurons aligned to the onset of the lick during perceptual licking. **o**, Changes in lick latency (black solid, 26 sessions), network activity from neighboring neurons in the ACC (gray, n = 450) during a session according to the time elapsed. **p**, Same as **o**, but for spontaneous licking rates (26 sessions). n.s. (not significant), \**p* < 0.05, \*\**p* < 0.01, \*\*\**p* < 0.001; Wilcoxon signed-rank test. Error bars show ± SEM.

Interestingly, we found that network activity in the population of ACC neurons that were not S_Inc_ or M_Dec_ neurons (Others) showed a significant reduction in the activity prior to spontaneous and perceptual licking but not during miss trials (Fig. 6j-l). Our data further showed that the level of suppression in these neurons significantly correlated with response times during perceptual licking: the lower the activity, the faster the licking (Fig. 6m,n). These results indicate that the decrease in the activity of neighboring neurons precedes S_Inc_-to-M_Dec_ suppression; network activity likely supports the sustained activity of M_Dec_ neurons. In other words, reduced network excitation is important for gating the premotor suppression of M_Dec_ neurons. We also found that network activity increased gradually along with lick latency across a session (Fig. 6o,p). Therefore, it is possible that network activity may be modulated along with changes in the internal states, such as the level of motivation or task engagement.

### Inactivation and measurement of ACC activity during the stop signal task

Because we observed that the sustained activity of ACC neurons is required for the suppression of impulsive licking (Fig. 1h-k), we wondered whether ACC neurons also play important roles in action cancellation during the stop signal task^25^. During the task, we presented auditory beeps as stop signals at randomly delayed times after the visual stimuli in a small portion of Go trials (Fig. 7a). Mice learned both to withhold licking prior to visual stimuli (levels of spontaneous licking were lower) and to stop licking after the stop signals (Fig. 7b, left). Muscimol-inactivation of ACC significantly increased both “waiting” impulsivity (Hit_IL_ trials increased) and “stopping” impulsivity (unsuccessful stop trials increased) (Fig. 7b-d). When we compared the degree of impairment in waiting and stopping licks (impairment index, MUS/PBS), ACC inactivation induced more impairments in waiting than in stopping (Fig. 7e). We next measured neural activity in the ACC during the stop signal task to analyze activity changes of ACC S_Inc_, M_Dec_ and other neurons when licking stopped (Fig. 7f,g). Notably, the activity of both ACC M_Dec_ and neighboring neurons (though not S_Inc_ neurons) increased significantly when mice stopped licking after the auditory cue (Fig. 7g). These results suggest that gating M_Dec_ suppression in the ACC is important for inhibiting licking, both on-going as well as spontaneous, during the task.

**Fig. 7.**
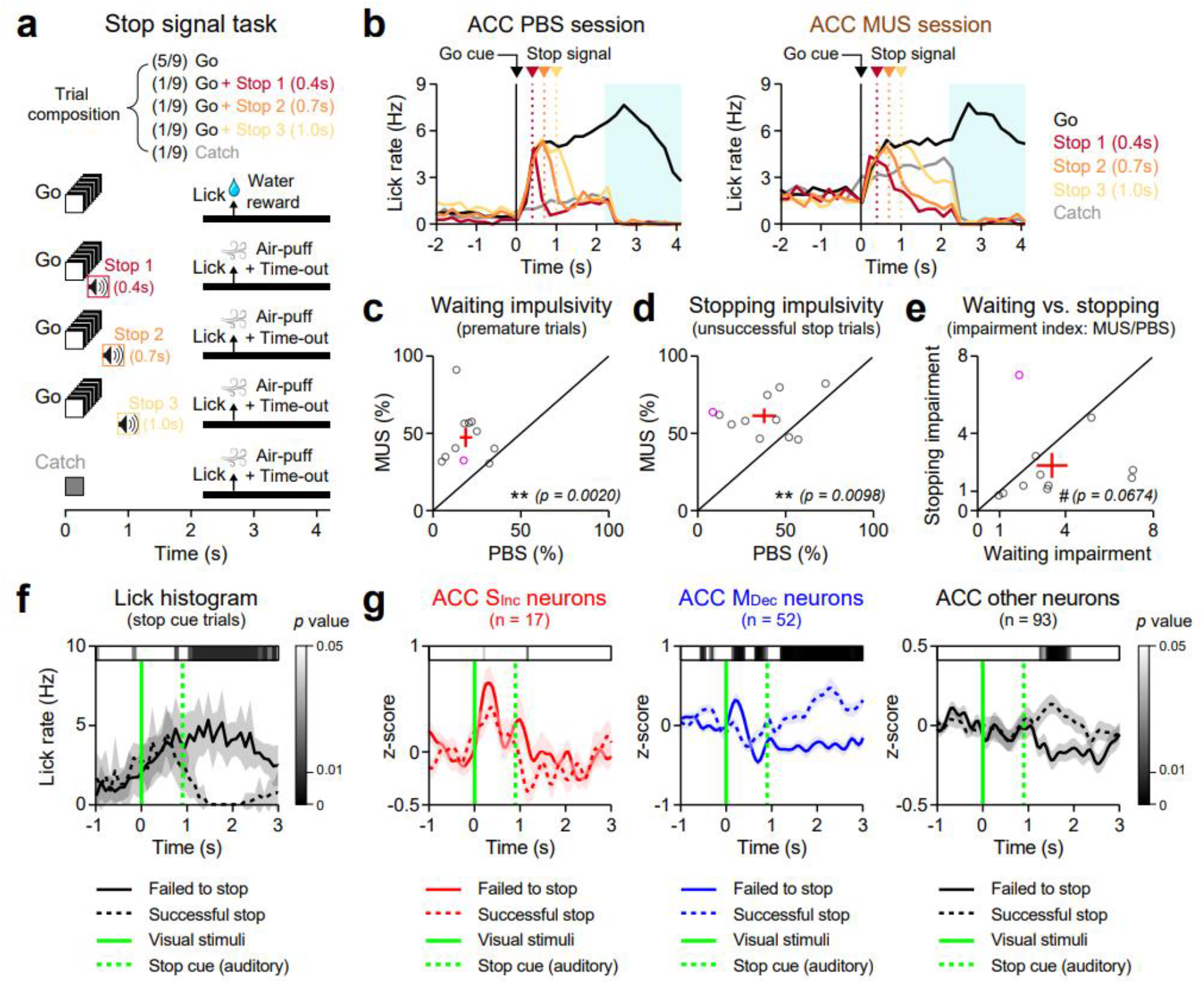
Role of the ACC in inhibitory control during the stop signal task. **a**, Schematic illustration of the stop signal task. **b**, Lick histograms of example mouse performing stop signal task after the injection of PBS (left) or muscimol (right; MUS) in the ACC during Go trials (black), Stop trials (red, 0.4 s delay; orange, 0.7 s delay; yellow, 1.0 s delay) and Catch trials (gray). **c-e**, MUS-induced impairments in task performance. % of premature licking trials (**c**, waiting impulsivity) and unsuccessful stop trials (**d**, stopping impulsivity) in mice injected with PBS or MUS. Gray circles, individual PBS-MUS sessions (n = 11, 7 mice); the magenta circle, the only session when MUS impaired stopping more than waiting; red crosses, means ± SEM. **e**, Degrees of impairments in waiting and stopping during the task. Note the tendency of increased waiting impulsivity more than stopping impulsivity after MUS injection into the ACC. **f**,**g**, Lick histograms (**f**) and neural activities (**g**) in the ACC S_Inc_ (left, red), M_Dec_ (center, blue), and other neurons (right, black) (G; z-scored) during unsuccessful (solid line) and successful stop trials (dotted line) averaged across 9 sessions of the stop signal task (5 mice). Shades, SEMs. Green vertical lines: solid, onset of visual stimuli; dotted, onset of stop auditory stimuli. Horizontal bars, color-coded *p*-values for significance difference between solid and dotted lines at each time bin (**f**, 200 ms; **g**, 50 ms). #*p* < 0.10, \*\**p* < 0.01; Wilcoxon signed-rank test.

Taken together, our data demonstrate that inhibitory inputs from S_Inc_ neurons and excitatory inputs from neighboring neurons are acting together to gate the suppression of M_Dec_ neurons, and such suppression releases both visually driven and spontaneous licking in task-performing mice at different times (Fig. 8).

**Fig. 8.**
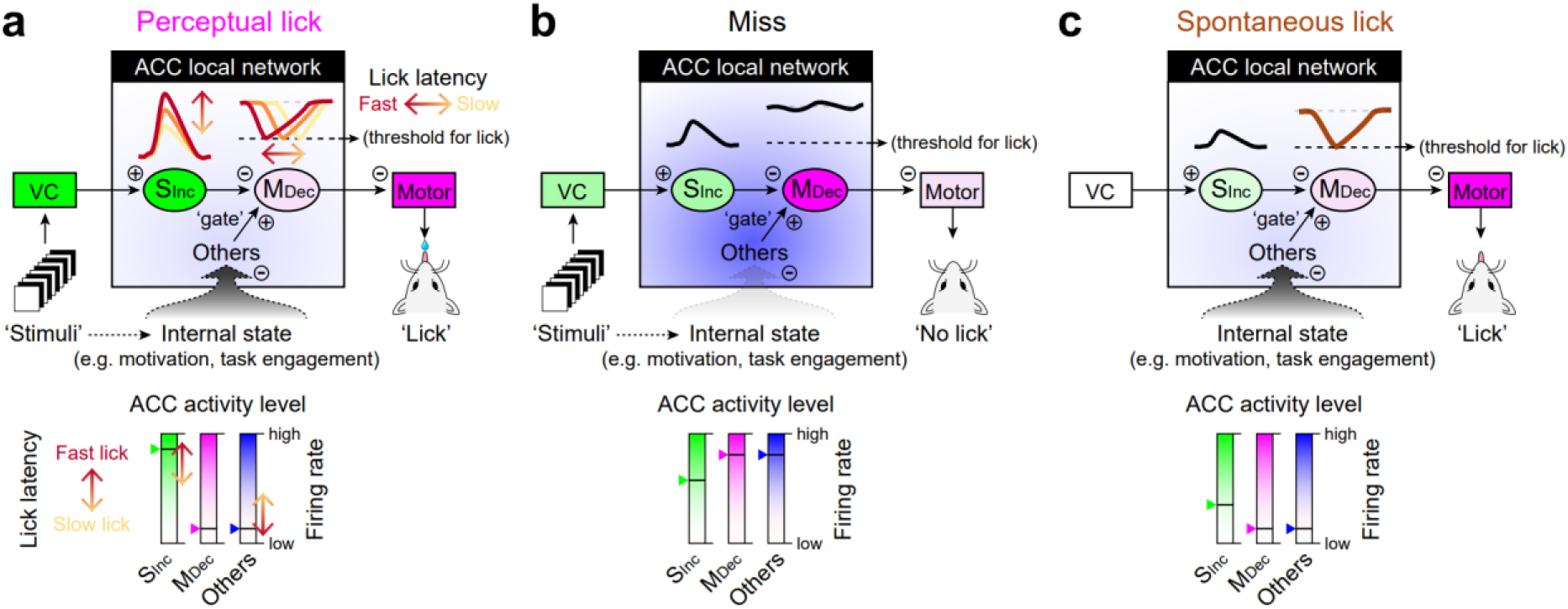
Schematic illustration of neural mechanisms underlying gated sensory-to-motor transformation in the ACC. **a**, A model of gated visual-to-motor suppression during ‘perceptual licking’ in the ACC. **b**,**c**, Illustration of ‘miss’ (**b**) and the ‘spontaneous licking’ (**c**) conditions in the same ACC network.

## Discussion

In this study, we found that the ACC receives visual inputs through S_Inc_ neurons and triggers motor action via the gated feedforward inhibition of the premotor activity in M_Dec_ neurons; the network activity in the ACC gates this sensory-to-motor suppression to release the licking response. Feedforward inhibition is a prominent feature of the bottom-up cortical processing^26,27^. Our data further demonstrate that the bottom-up inputs to the frontal cortex exert gated inhibition on the motor neurons (M_Dec_) that otherwise inhibit the lick responses. We propose that a balance between network excitation and sensory-induced inhibition determines when the M_Dec_ neurons are suppressed in the ACC, which in turn triggers the act of licking. Our data further support the idea that sustained activity in the ACC is important for the inhibitory control of licking responses in task-performing mice. In our study, M2 inactivation in mice did not impair their ability to perform visual tasks; other studies, in contrast, have reported that the motor cortical inactivation caused defects in perceptual decision-making in auditory or somatosensory tasks^5,28^. As shown by anatomical tracing of sensory inputs to the frontal cortex^16^, it is possible that the frontal cortex may be constructed with discrete areas that process perceptual decisions based on specific sensory information.

Our data showed that visual gains in S_Inc_ neurons in the ACC but not in the VC represent future reaction times. These results are in line with previous findings of transient activity in rat M2^5^ or visual neurons in monkey FEF^8^. Importantly, the gain modulation of S_Inc_ signals becomes more prominent in the ACC than in the VC. This result indicates that the sensory signal in the higher-order frontal cortex is more modulated by the task than in the lower sensory cortex, as shown in animals performing somatosensory tasks^29,30^. The visual gain in ACC S_Inc_ neurons might represent perceptual confidence in mice; cortico-cortical inputs, local circuitry within the ACC, and neuromodulators may work together to modulate visual representation in the ACC during a goal-directed perceptual task^31–33^. We further found that visual neurons in the ACC that receive direct synaptic inputs from the VC are largely FS inhibitory neurons. Feedforward inhibition is critical for balancing excitation and inhibition in cortical flows^34,35^. Our data further suggest that feedforward inhibition in the frontal cortex can be tuned by gain modulation in the visual responses of the FS neurons. As we observed this strong modulation only in the ACC, not in the V2M, it is possible that cell-type-specific neuromodulation in the frontal cortex may cause the specific modulation of FS S_Inc_ neurons in the ACC^36^. Interestingly, ACC S_Inc_ neurons show an increase in their activity even without visual stimuli prior to spontaneous licking. Whether this increase is also induced by the same mechanism that causes the gain modulation in S_Inc_ neurons needs future investigation.

The premotor suppression of M_Dec_ neurons in the ACC showed ramping-down property reaching a threshold at which mice initiated vision-guided licking. This ramping property is similar to other motor neurons in the cortex that represent trial-by-trial variations in response times of task-performing animals^5,7,37–39^. Similar to some FEF neurons^8,40^, M_Dec_ neurons showed a discrete rather than a continuous ramping activity. This indicates that feedforward inhibition from S_Inc_ neurons accumulates and overcomes the recurrent excitation to trigger the ramping-down activity in M_Dec_ neurons. Future experiments will be required to understand the synaptic microcircuits that integrate inhibitory inputs from the SInc neurons and excitatory inputs from the network in M_Dec_ neurons in the ACC. Defining cell types and projections of ACC M_Dec_ neurons will be important to understand a complete circuit diagram that governs sensory-evoked action execution. We did not observe any direct projection from the ACC to the anterior lateral motor cortex that is known to induce licking in mice^41^. Rather, ACC sends signals to the basal ganglia as well as the superior colliculus and the lateral hypothalamus^23,42,43^. The activity in these projections works as a default “licking withholder,” and sensory inputs to the ACC brake the activity to initiate the licking responses. In this process, M_Dec_ neurons determine when to release a stereotyped licking response similar to the fixation neurons in the primate FEF that produce eye movements^44^. Our finding accounts most fully for the function of the frontal cortex, which maintains both response inhibition before the sensory stimulus and decision-making upon its arrival. Consistent with this idea, hypoactivity in the frontal cortex has been observed in both animal models and human patients with psychiatric disorders that show an impairment of inhibitory control^45^, such as attention-deficit/hyperactivity disorder^46^, drug addiction^47,48^, and schizophrenia^49^. Our data also support that the inhibitory control of action is a prior cognitive function of the brain to perform goal-directed perceptual behaviors^50^.

The dual function of the ACC is implemented by maintaining distinct sensory and motor processors that are embedded in a network of neighboring neurons. The sensory processor consists largely of fast-spiking inhibitory neurons that can inhibit the motor processor and so trigger actions. The sensory-to-motor suppression is gated by the network activity that nearby neurons construct, as the inhibition occurred only when network activity was low. That is, the network activity from neighboring neurons attenuated the suppression of motor neurons even though the sensory signal was prominent in miss trials. In the new model that our data propose, the frontal cortex has a dual function, inhibitory control and perceptual decision making, by gating feedforward sensory-to-motor inhibition: it balances response inhibition and disinhibition. Neuromodulatory systems may be involved in regulating the level of network activity during the task^32,33^. Future studies are required to understand better the role of neuromodulatory inputs to the ACC in gating sensory-to-motor transformation.

## Methods

### Animals and surgery

All experimental procedures were approved by the KAIST Institutional Animal Care and Use Committee (IACUC-14-145). Animals were maintained *ad libitum* under light (8 am - 8 pm) and dark cycle (8 pm - 8 am) conditions, and housed under single animal per cage conditions from the first to the last day of experiments. We used C57BL/6 wild-type mice and *ROSA26^Ai^*^14^ Cre-dependent tdTomato reporter mice (Ai14, stock no. 007914 from the Jackson Laboratory) of both sexes (2 - 6 months old) and denoted all the numbers of mice used in each experimental condition in the Fig. legends and method details.

Animals were anesthetized with avertin (2, 2, 2-Tribromoethanol (Sigma-Aldrich), 125-250 mg/kg, intraperitoneally) and an additional 0.5 ∼ 1% isoflurane in O_2_ for stable maintenance of anesthetized status during surgical procedures. Body temperature was maintained at 37°C through a heating pad connected to a temperature feedback system (CWE Inc.). For retrograde tracing, we injected ∼0.5 μl of retrograde adeno-associated virus expressing tdTomato^51^ (retroAAV-tdTomato, Addgene Cat# 59462-AAVrg) into the right ACC (bregma +1.2 mm, lateral 0.3 mm, depth 1.2 mm) from wild-type mice using Nanoliter 2010 injector (WPI). For anterograde trans-synaptic and retrograde tracing, we injected ∼0.5 μl of anterograde trans-synaptic virus (AAV1-hsyn-Cre, Addgene Cat# 105553-AAV1) into the right DMS (bregma 0 mm, lateral 1.5 mm, depth 2.2 mm) and retroAAV-GFP (Addgene Cat# 37825-AAVrg) into right V2M (bregma -2.3 mm, lateral 1.5 mm, depth 0.7 mm) from Ai14 mice. To prepare mice for behavior and recording, the skin on the head was incised and remaining connective tissues were removed. We marked ACC, V1 (bregma −3.5 mm, lateral 2.5 mm, depth 0.7 mm), V2M, S1 (bregma -0.9 mm, lateral 3.0 mm, depth 0.8 mm), M2 (bregma +1.2 mm, lateral 1.2 mm, depth 1.0 mm), or DMS on the skull of the right hemisphere in order to carry out pharmacological inactivation experiments or *in vivo* extracellular recordings during the task. After marking the regions of interest (ROIs) and/or injecting virus, we attached a custom-designed head-plate onto the skull with small screws (Small Parts) and dental cement (Lang Dental). For optogenetic experiments, we injected virus on the target brain areas during the head-plate implant surgery. We continuously monitored the health of the head-plate implanted mice and started behavioral training one week after recovery.

## Behaviors

During behavioral tasks, mice were head-fixed and located in a soundproof behavior box. For presenting visual stimuli, a gamma-corrected LCD monitor was placed 10 cm away from the left eye of mice. Default visual luminance of the monitor was 60 lux (gray screen) during the task. Licking was detected by either a transistor-based lickometer, which senses the physical touch of the mouse tongue to a lick port (for behavioral training and pharmacological inactivation), or infrared beam-break sensors, which detect the tongue moving to the lick port (for electrophysiological recording to avoid electrical noise). We delivered water into the lick port via polyurethane tubing by gravity flow under the control of a solenoid valve (EV-2-24, Clippard). Task was operated by Presentation (Neurobehavioral Systems) with custom-written scenarios, custom-made electric circuits and USB data acquisition devices (USB-201, Measurement Computing).

### Visual detection task

In this task, we trained mice to detect visual stimuli under a simple GNG paradigm, as previously described^52^. Training proceeded as follows: reward habituation, conditioning, detection training, and detection tests with multiple luminance of visual stimuli. For reward habituation (3 ∼ 5 sessions), thirsty mice could get ∼3 μl of water whenever they licked the lick port. We moved to the next phase if mice executed over 1000 licking trials in 3 consecutive sessions. In the conditioning step (4 ∼ 5 sessions), a full field flashing light stimuli (10 Hz for 500 ms, five times repetition of 120 lux (50 ms) and 0 lux (50 ms) screen) was presented, and mice could get ∼3 μl of water reward for 4 s (Hit) if they licked during a 2 s response window (from 1 ∼ 3 s after stimuli onset). Even though they did not lick during the response window (Miss), a water reward was still given at the end of the response window. Mice exceeding 300 hits in 3 consecutive days were advanced to the next step. During the detection step (∼10 sessions), background (BG; no stimuli, default luminance (gray, 60 lux)) trials were randomly interleaved with Go trials (presenting equal stimuli used in conditioning), and mice were punished with a mild air-puff (300 ∼ 500ms) and a longer (8 s) inter-trial interval (ITI) if licks were detected during the response window of background trials (false alarm, FA). Mice were neither rewarded nor punished for miss trials and correct rejection trials (CR, no lick for response window of background trials); instead we just presented 3 s ITI after the end of the response window. Since mice started licking even before the onset of visual stimuli in some Go trials, we additionally divided hit trials into ‘perceptual lick’ hit trials (Hit_PL,_ no licking event during the 2 s period before onset of visual stimuli) and ‘impulsive lick’ hit trials (Hit_IL,_ at least one licking event during 2 s period before the onset of visual stimuli). Tasks were automatically ended when the number of consecutive miss trials exceeded 10. We calculated Hit_PL_, Hit_IL_, Miss, FA, CR rates and correct rate as follows:

*Hit_PL_ rate (%) = ((the number of Hit_PL_ trials) / (the number of Go trials)) x 100*
*Hit_IL_ rate (%) = ((the number of Hit_IL_ trials) / (the number of Go trials)) x 100*
*Miss rate (%) = ((the number of Miss trials) / (the number of Go trials)) x 100*
*FA rate (%) = ((the number of FA trials) / (the number of BG trials))) x 100*
*CR rate (%) = ((the number of CR trials) / (the number of BG trials))) x 100*
*Correct rate (%) = ((the number of Hit and CR trials) / (the number of all trials)) x 100*

We considered mice as experts in performing the task if they showed more than 70% correct rates over 3 consecutive days, and most mice reached the expert status within 10 sessions of the detection task. At the final step, we presented three weaker luminance (25%, 50% and 75% compared to luminance of Go stimuli (120 and 0 lux)), randomly interleaved with Go and background trials at different probabilities (3/8, 2/8, 1/8, 1/8, and 1/8 for Go, BG, 25%, 50%, and 75% luminance trials, respectively). Stimuli structures in the test trials with weaker visual stimuli were equal to Go stimuli, but we did not give water rewards during the response window.

### Stop signal task

If in the detection training step, mice learned the visual detection task at 70% correction rates, we presented the auditory stop cue (10 Hz pure tone, 70 dB) after the Go visual stimuli at various stop signal delays (SSDs; 0.4 s, 0.7 s, and 1.0 s) (stop trials; 1/9 of total trials for each type of SSDs). We gave an air-puff punishment if mice did not stop licking 0.5 s after the auditory cue and counted such a stop trial as unsuccessful. Similar to Hit_IL_ trials in the detection task, if mice started licking within 2 s period prior to the visual stimuli among hit trials, these trials were counted as premature. We quantified the percentage of premature trials among total hit trials and the percentage of unsuccessful stop trials among stop trials as indices for waiting impulsivity and stopping impulsivity, respectively. We also punished mice if they licked in the catch trials (background stimuli; 1/9 of total trials). To compare the degree of waiting impulsivity with that of stopping impulsivity, we calculated impairment indices by dividing the waiting and stopping impulsivity (% of premature and unsuccessful stop trials) with muscimol injection (MUS) by those with PBS injection (PBS).

#### Pharmacological inactivation

For pharmacological inactivation, we unilaterally injected 0.23 μl of either GABA_A_ receptor agonist muscimol (MUS, 1 μg/μl, dissolved in PBS, Sigma-Aldrich) or PBS into the right ACC, V1, V2M, M2, and S1. Twenty minutes after MUS or PBS injection, we measured the performance of mice during the visual detection task (ACC, 14 mice; V1, 10 mice; V2M, 5 mice; M2, 4 mice; S1, 11 mice), and during the stop signal task (ACC, 7 mice). In order to compare the effects of MUS with PBS in the same animal, we trained the same mice across multiple sessions in the following order: pre (no injection), MUS injection, PBS injection, Fluorophore-conjugated MUS (FCM) injection.

### *In vivo* electrophysiological recordings

#### Neural activity recording during task

We performed *in vivo* extracellular recordings in ACC, V1, V2M, M2, or DMS from trained mice performing behavioral tasks (ACC, V1, V2M, M2, or DMS for the visual detection task; ACC for the stop signal task). Before the initiation of the behavioral task, a small craniotomy (diameter ∼0.5 mm) was made over the target area, into which we slowly inserted a single shank silicon electrode with 32 recording sites (A1x32-Poly3-10mm-50-177-CM32, Neuronexus) using a micro-drive motorized manipulator (Siskiyou). After waiting 30 minutes for the silicon electrode to stabilize at the targeting site, we started the behavior task and concurrently recorded neuronal activity, filtered between 500 ∼ 5000 Hz at a 30 kHz sampling rate and amplified by a miniature digital head-stage (CerePlex μ, Blackrock Microsystems), using a data acquisition system (CerePlex Direct, Blackrock Microsystems) for offline single unit isolation. To confirm recording sites and for post hoc histology analysis, we coated a silicon electrode with liphophilic dye (1,1’-dioctadecyl-3,3,30 3’-tetramethylindocarbocyanine perchlorate (DiI) stain, Invitrogen).

#### Neural activity recording from untrained mice

After mice finished reward habituation step (see Behavior in Methods), we recorded neural activity in the ACC of untrained mice using a silicon electrode. To measure both sensory and motor signals, we either presented visual stimuli or gave water with random ITIs (5 ∼ 8 s) during recording. We stopped recording after collecting at least 30 visual trials and 30 motor (spontaneous licking) trials. All the recording steps were implemented in the same manner as described above.

#### Opto-tagging recording

To measure the activity of V2M-recipient ACC (ACC_←V2M_) neurons, we expressed ChR2 in ACC_←V2M_ neurons by injecting ∼0.5 μl of AAV1-hSyn-Cre into the V2M and ∼0.5 μl of AAV2-EF1α-DIO-hChR2-mCherry (UNC Vector Core) into the ACC of the right hemisphere. After mice learned the task, we inserted an opto-silicon electrode (A1x32-Poly3-10mm-50-177-OCM32LP (105 μm core diameter with 0.22 NA), Neuronexus), connected to a blue laser (473 nm, ∼1 mW, Shanghai Laser & Optics Century), into the right ACC. Light stimulation was controlled by a pulse stimulator (A-M Systems). To identify ACC_←V2M_ neurons (opto-tagged), we delivered 1Hz, 200 light pulses (10 ms) before task initiation, and continuously recorded neural activity during task performance.

#### Measuring effects of muscimol on neural activity

To confirm that MUS could disrupt neural activity, we performed *in vivo* extracellular recording in ACC under head-fixed status. Using silicon electrode, we firstly measured about 10 minutes of baseline activity, and applied 3 μl of PBS onto the recording site to check effects of PBS on neural activity. About 40 minutes later, we applied 3 μl of MUS onto the same site. We terminated recording 40 minutes after MUS application. Data acquisition was carried out with the same way as described above.

### Optogenetic manipulation of neurons during the task

We measured effects of photostimulation on ACC_←V2M_ neurons (soma stimulation, ChR2), ACC-projecting V2M axons (axon stimulation, ChR2), and control groups (eYFP or no virus injection) by simultaneous measuring the behavioral performance of mice and the neural activity. To label ACC-projecting V2M axons, we injected ∼0.5 μl of AAV2-CaMKIIα-hChR2-eYFP (UNC Vector Core) into the V2M of the right hemisphere. During the task, we delivered five light pulses (473 nm, ∼1 mW; 50 ms per pulse) at the same temporal structure with the actual visual stimuli, in random trials interleaved with the actual visual stimulation trials (Go (100%) or 50% luminance) and the BG trials. We conducted opto-tagging recordings and photostimulation experiments with different groups of mice (see Supplementary Table 1). Licking rates (Hz) were calculated from lick events during 2 s response window in each trial, and lick probability (%) was defined as the fraction of trials when mice showed licking during 2 s response window across trial types.

### Histology and imaging

We performed histology experiments for anatomical circuit tracings, confirmation of targeting sites after pharmacological inactivation experiments, and electrode track identification after *in vivo* recordings. Mice were deeply anesthetized with avertin and transcardially perfused with PBS followed by 4% paraformaldehyde (wt/vol in PBS). Brain samples were isolated and post-fixed for 4 hours and put in filtered 30% sucrose solution (wt/vol in PBS) until they sank completely. Afterward, the samples were embedded with optimal cutting temperature (OCT) medium and rapidly frozen at −80°C. Frozen brain samples were sectioned into 40 μm thickness of coronal slices using a cryostat (Leica). We washed brain sections 3 times for 10 minutes each with 0.1 M phosphate buffer (PB), and mounted them with 4′,6-diamidino-2-phenylindole (DAPI) containing mounting medium (Vector Labs). To perform immunostaining of the parvalbumin (PV) in brain samples labeled with eGFP in ACC_←V2M_ neurons, we permeabilized the brain sections for 30 min with 0.1% Triton X-100 in 0.1 M PB solution and incubated them in the blocking solution (2% normal donkey serum (NDS) in 0.1 M PB) for 2 hours at room temperature (RT). We then treated the sections with rabbit anti-PV antibodies (1:500 dilution in 2% NDS; Swant) for 48 hrs at 4 °C. We next washed the sections 3 times with PB buffer and treated the secondary antibodies (1:500 dilution in 0.1 M PB; Alexa Flour 594 donkey anti-rabbit IgG, Thermo Fisher Scientific) for 2 hours at RT. After that, we washed the sections with PB buffer 3 times again and mounted them with DAPI solution. Fluorescent images of brain sections were taken by slide scanner (Zeiss Axio Scan.Z1). For ACC retrograde tracing, we automatically matched fluorescent images of coronal sections with Allen Reference Atlas (ARA) and automatically quantified fluorescence-labeled cells using a recently developed automatic 3-D mapping program^53^.

### Video monitoring of orofacial movements of mice performing the task

We monitored orofacial movements of task-performing animals with a camera (G3-GM11-M2020-Genie, Teledyne DALSA) and lens (TEC-M55 2/3” 55mm telecentric lens, C-Mount) at a high resolution (1200 x 800 pixels) by illuminating left side of the animal face with infrared light (850 nm, S-IR5850, Skycares). We recorded images at 30 Hz via Image Acquisition Toolbox provided by MATLAB.

### Neural activity analysis

#### Spike sorting for single unit isolation

Spike sorting was executed by separating 32 channels on the silicon electrode into eight groups (four proximate channels per group) and sorting distinct spike clusters based on three principal components displayed in Klusters software^54^. We checked that the auto-correlogram of each cluster had a 2 ms refractory period in order to rule out the possibility of multi-unit contaminations.

#### Classification of task-relevant responses

To examine the responsiveness of ACC, V1, V2M, M2, and DMS neurons to visual stimuli in task-performing mice, we calculated the firing rates (FRs) of individual neurons before (−0.5 ∼ 0 s) and after (0 ∼ 0.5 s) visual stimuli onset of hit trials. To rule out contamination to sensory signals by licking behavior *per se*, we used Hit_PL_ trials in which visual stimuli did not overlap with the licking bout, from onset to offset. Each neuron was classified as increasingly (S_Inc_), decreasingly (S_Dec_), or non-responsive to sensory stimuli based on bootstrap analysis (*p* < 0.01, n = 5000). To check responsiveness related to licking behavior during the task, we first analyzed onset time of licking bouts during task, and confirmed that these events were composed of bouts, with short inter-lick intervals (ILIs) within a single bout (< 2 s) and long ILIs between bouts (> 2 s), consistent with previous literature^2^. Then, we separated licking into spontaneous (licking onset during background trials and ITIs) and perceptual (licking onset during Go trials) licking bouts, and compared the FR of the neurons during the pre-period (− 1.5 ∼ -0.5 s from spontaneous licking onset) to those during post-period (0 ∼ 1 s from spontaneous licking onset). Each neuron was classified as an increasingly (M_Inc_), decreasingly (M_Dec_) or non-responsive to the onset of licking events based on bootstrap analysis (*p* < 0.01, n = 5000).

#### PSTH, PETH, and correlation analysis

To plot peri-stimulus time histograms (PSTH) or peri-event time histograms (PETHs), we normalized (z-scored) neural activity by baseline activity (−2 ∼ 0 s from the onset of visual stimuli for PSTHs and −2.0 ∼ −0.5 s from the onset of licking for PETHs). Visual responses from different luminance were calculated from mean z-socred activity during the 0.5 s period after visual cue onset. We quantified absolute sensory (|z|, 0 ∼ 0.5 s after stimuli onset) and premotor (spontaneous and perceptual licking, |z|, −0.5 ∼ 0 s from the onset of licking) responsiveness for all activity in the total population of untrained and trained mice. To investigate the correlation between visual response and lick latency, all hit trials were categorized according to variation in lick latency and calculated mean firing rates during the presentation of visual cues (0.5 s), and averaged lick latency from different segments of hit trials. To investigate the linear relationship between visual response and lick latency, we analyzed the Pearson correlation coefficient (*r*) and its statistical significance (*p*) for all S_Inc_ and S_Dec_ neurons. We presented firing rate (spikes/s) for plotting example neuron and population data. To study correlation between motor-related activity and lick latency, we calculated the Pearson correlation coefficient (*r*) and its statistical significance (*p*) for all M_Inc_ and M_Dec_ neurons from every time bin. PSTHs in PETHs were smoothed by convolution with a Gaussian filter (σ = 50 ms) only for visualization purpose (*smoothdata*, built-in MATLAB function). To analysis difference of neural activity from test trials (spontaneous licking (lick onset), perceptual licking (either stimuli or lick onset), and miss (stimuli onset)) with baseline activity (no stimuli, no licking), we extracted baseline trials which have no stimuli and no licking during 6 s. We calculated a normalized rate, relative to the baseline activity, in the range between −1 and 1.

#### Opto-tagging recording and spike waveform analysis

To be identified as opto-tagged (directly light-activated), neurons had to satisfy the following two criteria^55,56^. First, the latency distribution (10 ms time window) of the first spike after the onset of light pulse had to be significantly reduced (*p* < 0.01) compared with that of no-stimulation condition by conducting the stimulus-associated spike latency test (SALT)^56^. Second, waveform correlation between light-driven and spontaneous spikes must be more than 0.95. For spike waveform analysis, we classified the putative cell-type of recorded neurons based on spike width (time between peak and trough of the spike, μs), as previously described^41^. We considered recorded neurons as FS when their spike width was below 350 μs, and those as regular-spiking (RS) when their spike width was above 450 μs.

#### Prediction of lick latency and statistical test for identifying significant motor neurons

To predict lick latency M_Inc_ and M_Dec_ neurons from ACC, V1, and M2, we calculated the time required to cross threshold from distinct PSTHs of neurons with different lick latency, similar to a previous study ^5^. We first defined ‘threshold 1’ as the highest minimum neural activity (for M_Dec_) or the lowest maximum (for M_Inc_) for distinct PSTHs during 0 ∼ 3 s after stimuli onset, overlaid a horizontal line (threshold 1 as a y-intercept), and measured the time required to cross threshold from each PSTH during 0 ∼ 3 s after stimuli onset. Next, we defined ‘threshold 10’ as the lowest maximum neural activity (for M_Dec_) or the highest minimum (for M_Inc_) from distinct PSTHs during a baseline period (−2 ∼ 0 s prior to stimuli onset), overlaid a horizontal line (threshold 10 as an y-intercept), and measured the time required to cross threshold from each PSTH during 0 ∼ 3 s after stimuli onset. Additionally, we defined eight intermediate thresholds equally spaced between thresholds 1 and 10, and calculated the time required to cross thresholds in the same manner. Then, we carried out the Pearson correlation analysis between actual lick latency and predicted lick latency. To identify significant motor neurons, we used the three criteria described previously^5^. Briefly, (1) at least 2 of the 10 thresholds need to show significant correlations between lick latency and the time required to cross the threshold, (2) the time required to cross those significant thresholds need to precede lick onset time (a.k.a. positive prediction time), and (3) the regression slope for the time required to cross at least one of the significant thresholds against the lick time is close to unity (0.8 ∼ 1.2).

### Analysis of pupil size and video motion energy

We calculated pupil size of the left eye of the task-performing mice with a custom-written MATLAB code as follows: 1) drawing a ROI that fully covers the eye and the eyelid, 2) finding an optimal gray-scaled intensity as a threshold for defining the boundary of the pupil automatically, 3) converting the ROI image to a binary image by filling in each pixel with the black and white values according to the threshold (*imbinarize*, built-in MATLAB function), 4) filling holes within the boundary of the pupil to avoid any artifacts from measuring pupil area due to the light reflection on the cornea (*imfill*, built-in MATLAB function), 5) finding a convex hull including inside part of the pupil (*bwconvhull*, built-in MATLAB function), and 6) counting the number of pixels within the convex hull as the pupil size across frames. To measure orofacial movement, we quantified video motion energy (VME) of mouse orofacial images as previously reported^57^ using custom-written MATLAB codes. In brief, we drew ROIs covering individual behavioral parameters (either moving the nose or whiskers, or licking) and calculated video motion energy from each ROI through the following equation:

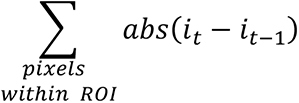

where *i_t_* is the gray-scaled intensity of the pixel on the frame *t*. To measure the behavioral changes during perceptual or spontaneous licking, we normalized (z-scored) both pupil size and video motion energy with the baseline values averaged for 2 s before licking.

### Statistical analysis

We used Excel (Microsoft) or custom-written codes in MATLAB (Mathworks) for data analysis. All data are displayed as mean ± standard error of the mean (SEM) unless noted otherwise. ‘N’ indicates the number of mice, ‘n’ the number of sessions or single units. Statistical differences were evaluated through bootstrap (classification of task response type only, *p* < 0.01), SALT (opto-tagging identification, p < 0.01), Wilcoxon signed-rank test (paired datasets), Wilcoxon rank sum test (un-paired datasets), or two-sample Kolmogorov-Smirnov test (nonparametric test for the equality between CDFs). Without bootstrap and SALT, all the other statistical significances of the datasets throughout the paper are shown as n.s., *, **, and *** indicate *p* ≥ 0.05, *p* < 0.05, *p* < 0.01, and *p* < 0.001, respectively.

## Acknowledgments

We thank Dr. Veit Stuphorn, Dr. Daeyeol Lee, Dr. Min Whan Jung, and all the other members of the Lee lab for helpful discussions. We also thank Emily Wheeler, Boston, for editorial assistance. This work was supported by grants to S.-H. L. from the Korean government through the National Research Foundation funded by the Ministry of Science and ICT (2018R1A4A1025616, 2017R1A2B3008270, 2017M3C7A1030798), the KAIST Global Singularity Program for 2020, and the ETRI grant (19ZS1500).

## Author contributions

J.-H.K. and S.-H.L. conceived and designed the experiments. J.-H.K. performed and organized all the experiments. D.-H.M. and E.J. performed some of the behavior training and histological experiments. I.C. established an infrared recording system. J.-H.K. and S.-H.L. performed data analysis and visualization. J.-H.K. and S.-H.L. wrote the paper.

## Competing interests

The authors declare no competing financial interests.

## Data and Code Availability

Licenses for Presentation, MATLAB, and Excel are commercially available, and the Klusters software can be freely downloaded at http://neurosuite.sourceforge.net/. All quantification methods used in the custom scripts are described in Method Details. Further requests for custom scripts and data used in this study can be directed to the corresponding author.

## Supplementary information

**Supplementary Fig. 1.**
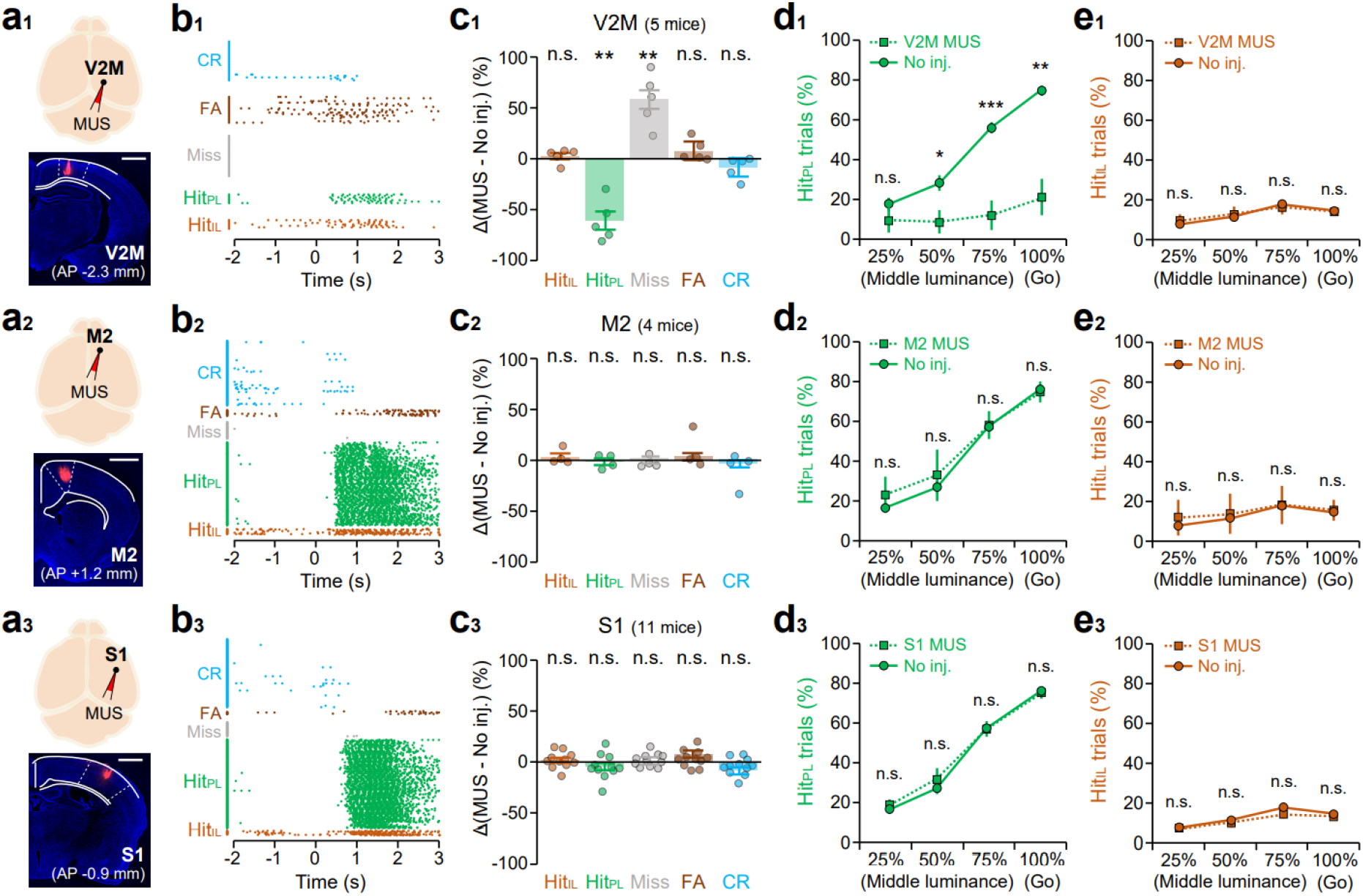
Effects of V2M, M2, and S1 inactivation on task performance. **a**, Schematic of MUS injection (top) and histological confirmation of MUS injection (bottom) into the V2M (**a**_1_), M2 (**a**_2_), and S1 (**a**_3_). Scale bars, 1 mm. **b**, Example lick raster plots of mice at each injection condition (V2M MUS (**b**_1_), M2 MUS (**b**_2_), and S1 MUS (**b**_3_)). **c**, % changes of Hit_IL_, Hit_PL_, Miss, FA, and CR rates by MUS injection compared with no injection (No inj.). **d**, % of Hit_PL_ trials across the trials with different levels of luminance. Dotted lines with squares, MUS injection; solid lines with circles, no injection controls. **e**, Same as **d**, but for Hit_IL_ trials. Color represents trial types. n.s. (not significant), \**p* < 0.05, \*\**p* < 0.01, \*\*\**p* < 0.001; Wilcoxon signed-rank test with Bonferroni correction. Error bars show ± SEM.

**Supplementary Fig. 2.**
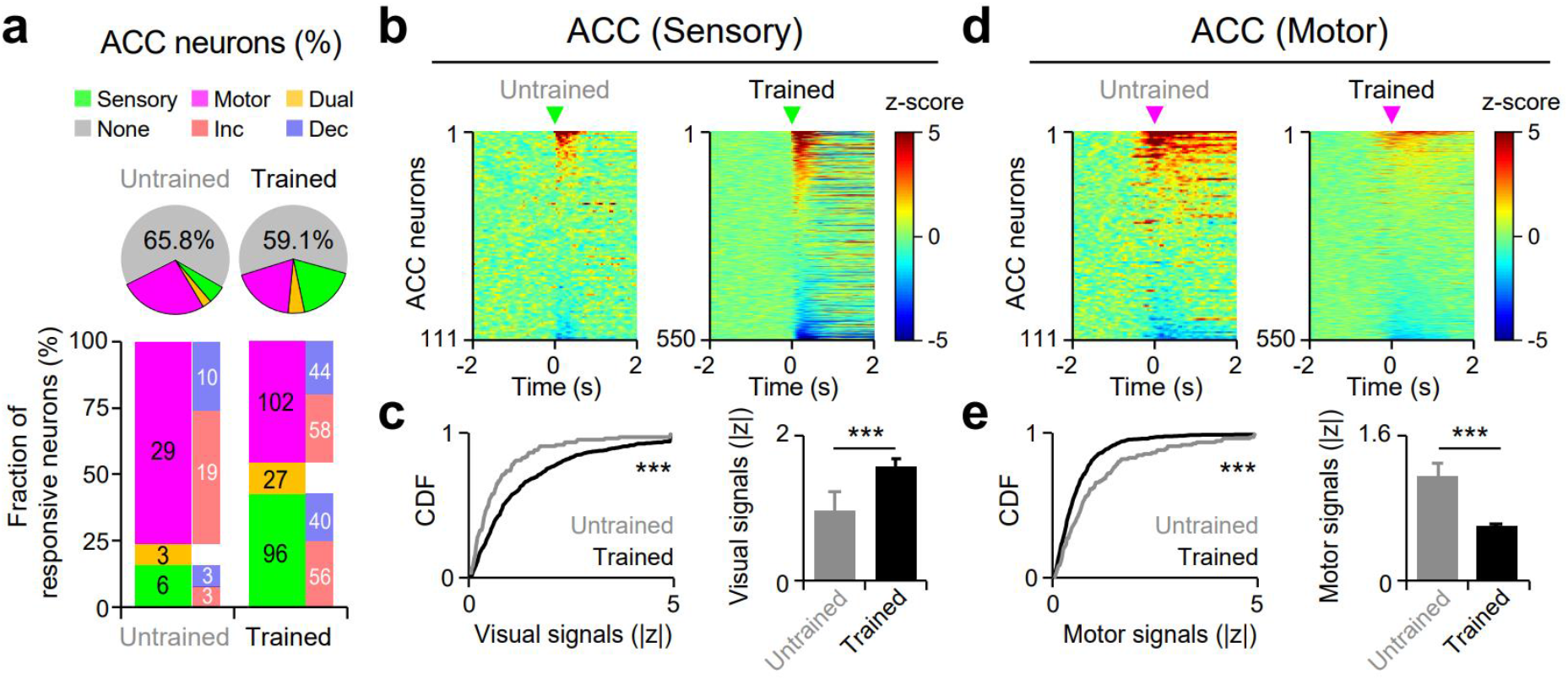
Learning-induced changes in ACC activities. **a**, Identification of task-relevant neurons in the ACC of untrained (left; n = 111, 3 mice) and trained mice (right; n = 550, 21 mice). **b**, z-scored population activity of ACC neurons showing visual responses from untrained (left) and trained mice (right). Green arrowheads indicate the time of stimuli onset. **c**, Cumulative distribution (left) and population average (right) of absolute visual signals (|z|, 0 ∼ 0.5 s after stimuli onset) in ACC neurons from untrained (gray) and trained mice (black). **d**,**e**, Same as **b**,**c**, but for motor signals of ACC neurons from untrained and trained mice. Absolute motor signals are calculated during −0.5 ∼ 0.5 s from lick onset. Magenta arrowheads indicate lick onset time. n.s. (not significant), \**p* < 0.05, \*\**p* < 0.01, \*\*\**p* < 0.001; Two-sample Kolmogorov-Smirnov test (CDFs in panel **c** and **e**) and Wilcoxon rank-sum test (bar graphs in panel **c**, **e**). Error bars, ± SEM.

**Supplementary Fig. 3.**
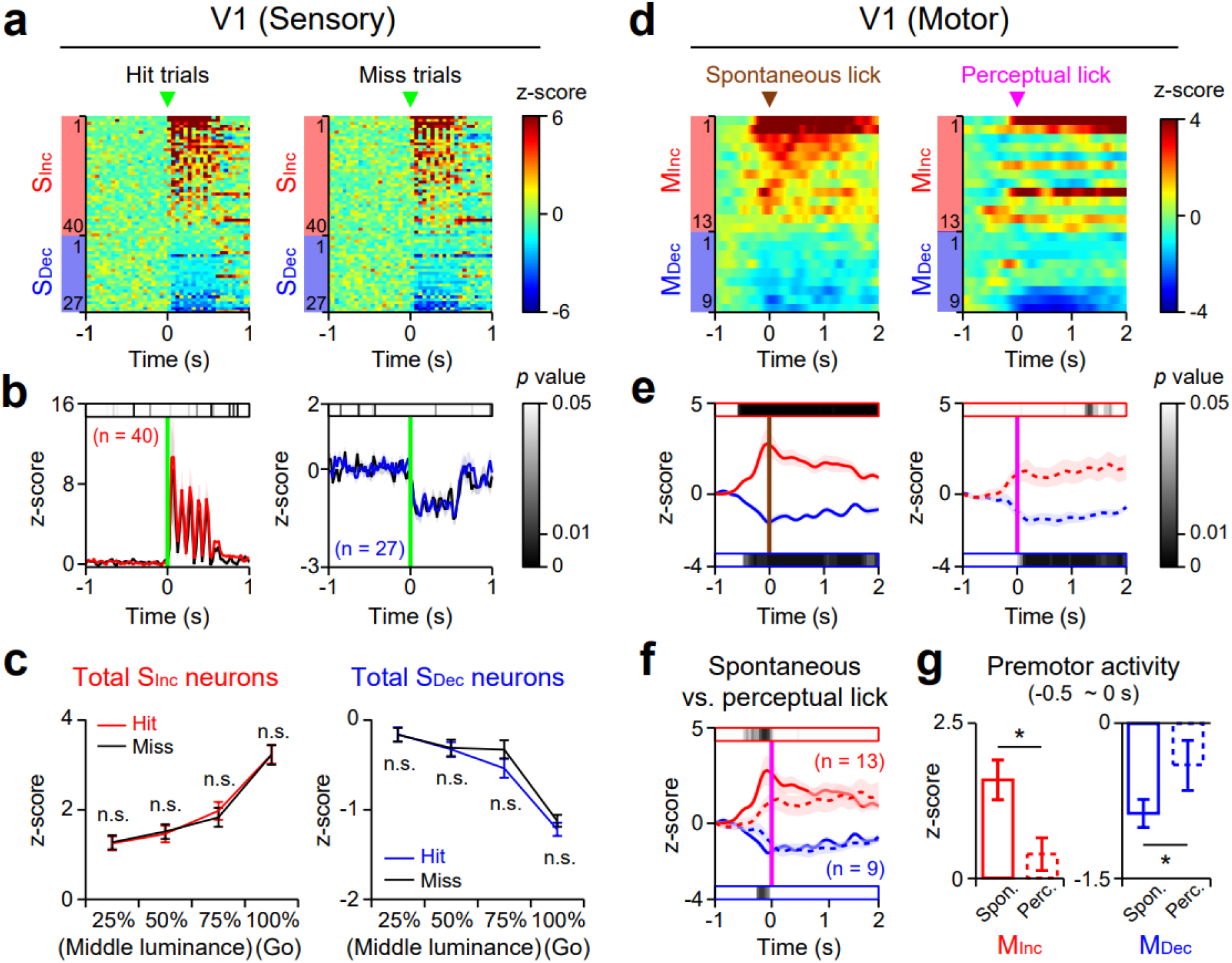
Sensory and motor signals in the population of V1 neurons. **a-c**, Same as Fig. 2**a-c**, but for V1 S_Inc_ (n = 40) and S_Dec_ neurons (n = 27). **d-g**, Same as Fig. 2**d-g**, but for V1 M_Inc_ (red, n = 13) and M_Dec_ neurons (blue, n = 9). n.s. (not significant), \**p* < 0.05; Wilcoxon signed-rank test. Error bars, ± SEM.

**Supplementary Fig. 4.**
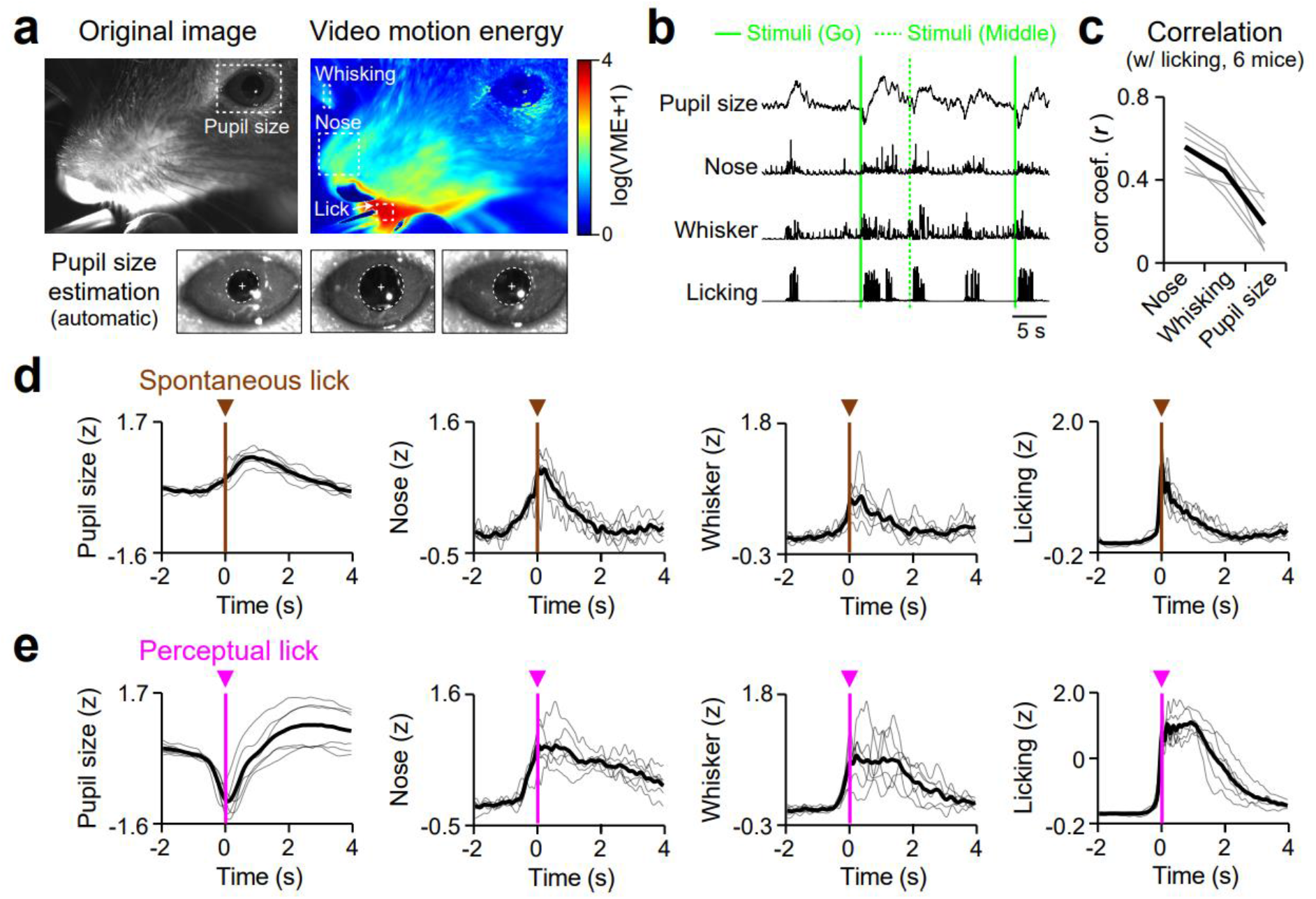
Analysis of orofacial movements in mice performing the visual detection task. **a**, Original image of the head of an example mouse taken by the infrared high-resolution camera (top left) and its video motion energy (VME, top right). Dotted rectangles, analyzed ROIs for pupil size estimation and VME of the orofacial movements (moving nose, whiskers, and licking). Bottom, the estimated pupil boundary of three representative images of the mouse eye (dotted ellipses). **b**, Fluctuation of pupil size and VME of three orofacial movements during visual detection task. Green solid lines and green dotted lines indicate the onset of Go and middle luminance stimuli, respectively. **c**, Pairwise correlation coefficient (*r*) between licking and other behavioral movements. Black line, mean; gray lines, individual mouse. Total 6 mice. **d**,**e**, Normalized (z-scored) fluctuations of four behavioral movements around the onset of spontaneous licking (**d**, magenta arrowheads and vertical lines) and the onset of perceptual licking (**e**, brown arrowheads and vertical lines). See also Supplementary Video 1.

**Supplementary Fig. 5.**
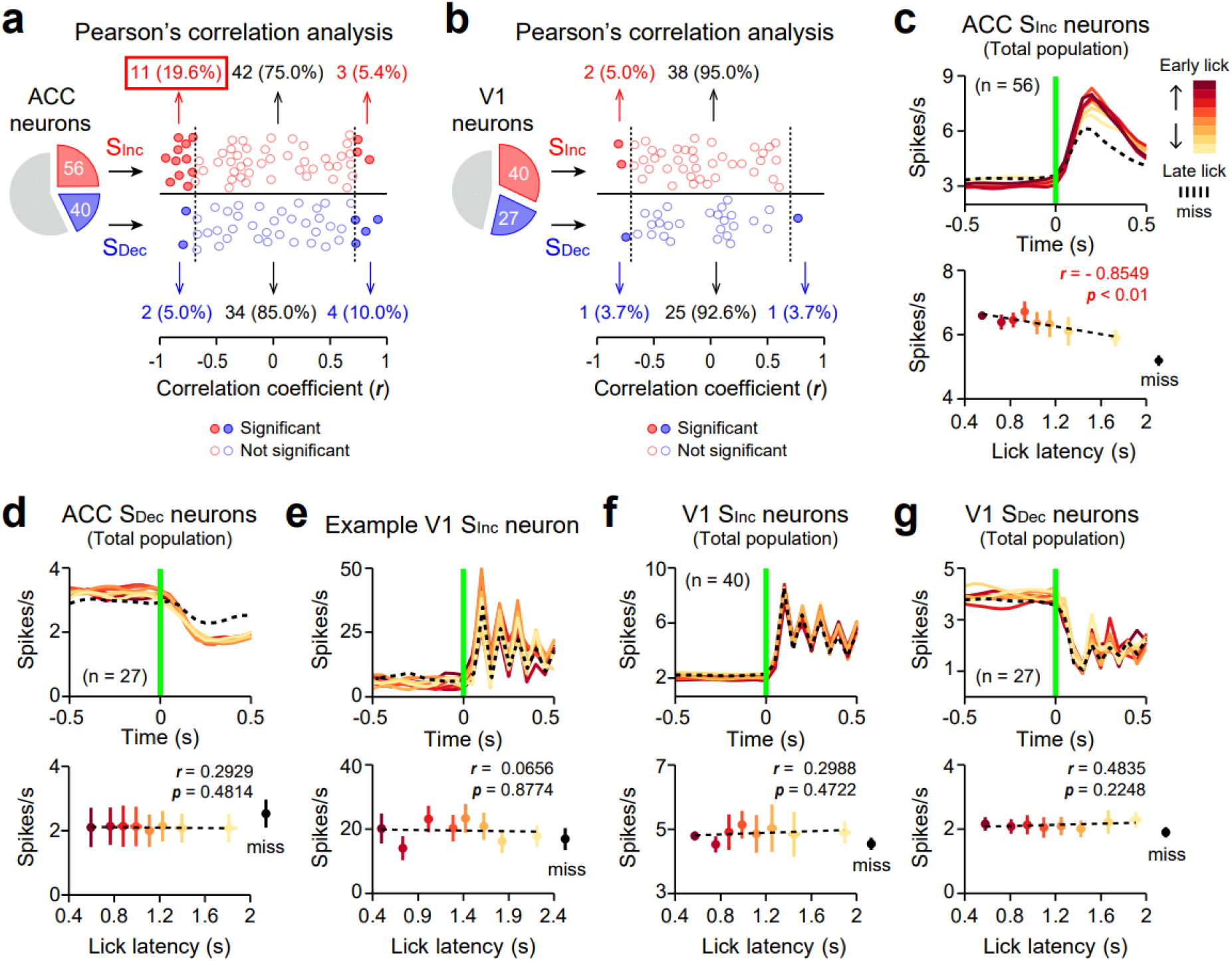
Correlation between licking response times (lick latency) and activities of S_Inc_ and S_Dec_ neurons in the ACC and the V1. **a**, Fraction of ACC S_Inc_ and S_Dec_ neurons that show significant correlation (Pearson correlation, *p* < 0.05) between visually evoked activities and lick latencies. Filled circles indicate neurons with significant correlation **b**, Same as **a**, but for V1 S_Inc_ and S_Dec_ neurons. **c-g**, Same as Fig. 2**j**, but for total population of ACC S_Inc_ neurons (**c**), total population of ACC S_Dec_ neurons (**d**), example V1 S_Inc_ neuron (**e**), total population of V1 S_Inc_ neurons (**f**), and total population of V1 S_Dec_ neurons (**g**).

**Supplementary Fig. 6.**
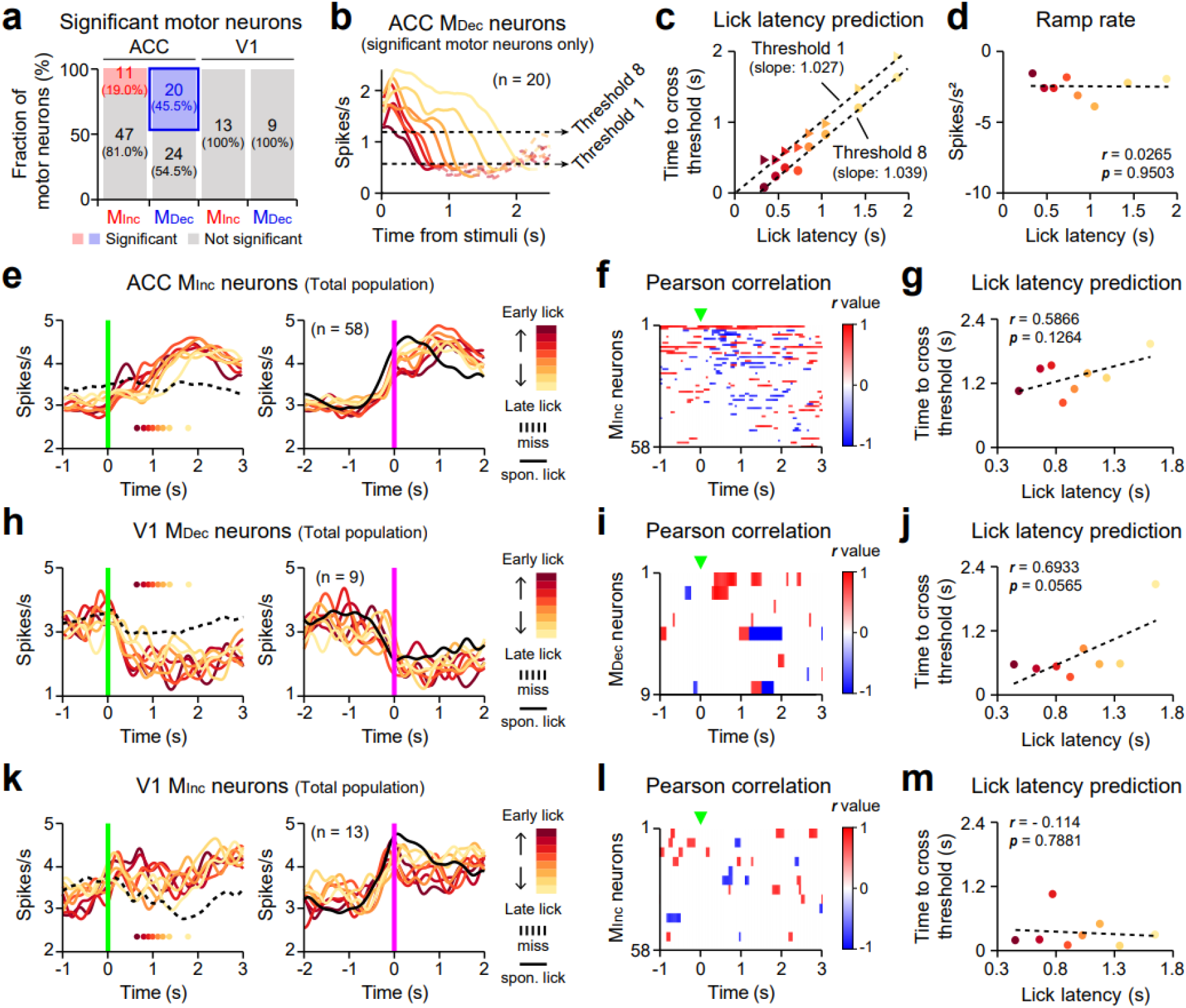
Correlation between licking response times (lick latency) and ramping activities of motor neurons in the ACC and the V1. **a**, Fraction of M_Inc_ and M_Dec_ neurons that show ramp-to-threshold premotor activity in ACC and V1. Red and blue, neurons that show significant correlation between time to cross the lower threshold and lick latencies. **b-d**, Color-coded PSTHs (**b**), scatter plots (**c**), and ramp rates (**d**) of ACC M_Dec_ neurons that show significant correlation in **a** (neurons in the blue box) across trials with early-to-late lick latencies. Note that time to cross thresholds (either 1 or 8) in ramping-down activity of ACC M_Dec_ neurons showed a linear relationship with lick latency at a slope near 1, while ramp rates (the slopes of ramping-down activity) were constant. **e-m**, Same as Fig. 2**l-n**, but for ACC M_Inc_ (**e-g**), V1 M_Dec_ (**h-j**) and M_Inc_ neurons (**k-m**).

**Supplementary Fig. 7.**
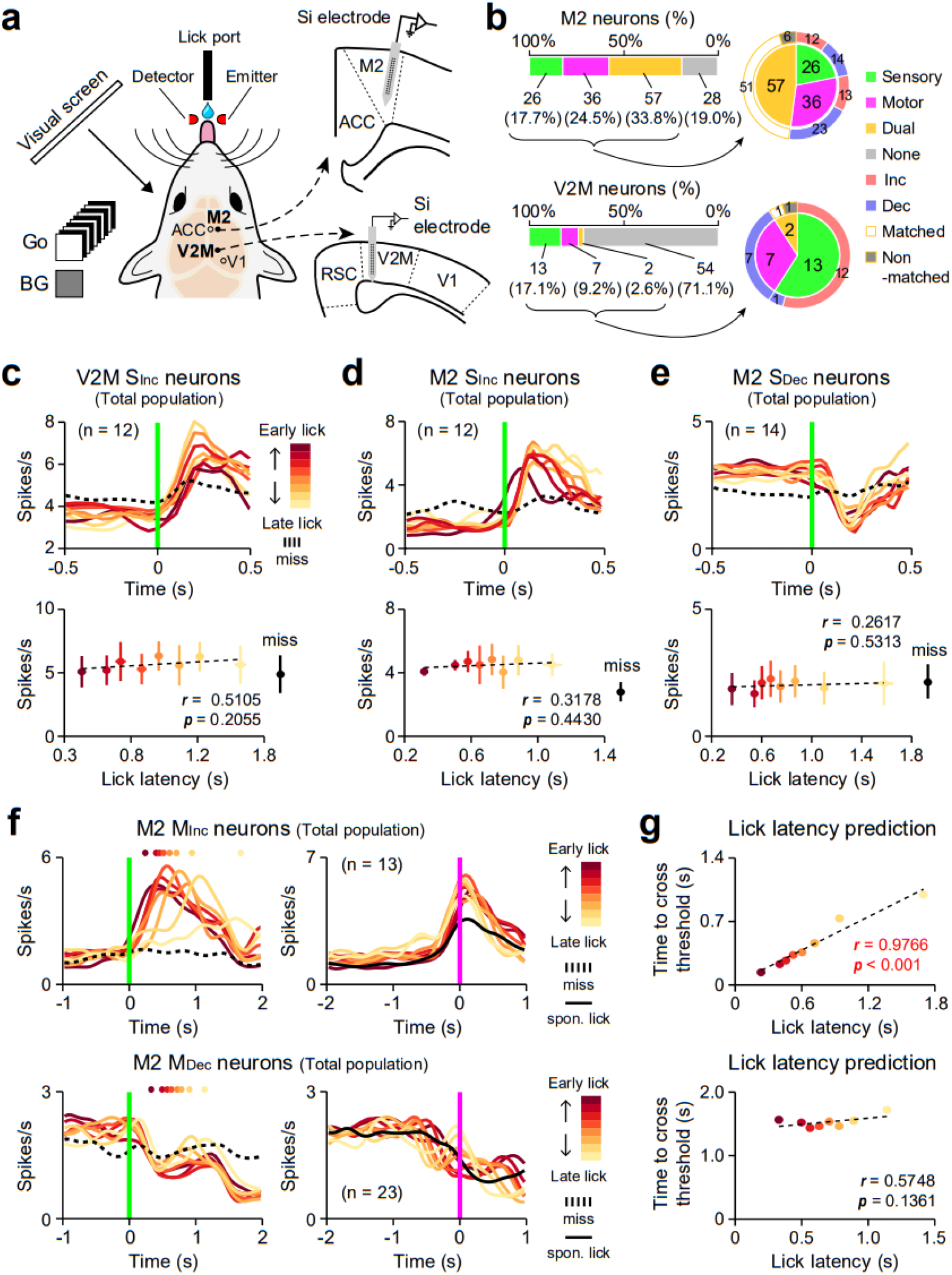
Task-related activity in V2M and M2 neurons. **a**, Schematic of *in vivo* multichannel recording in the V2M or M2 of mice performing the visual detection task. **b**, Categorization of neuronal types in the M2 (top; n = 147, 4 mice) and the V2M (bottom; n = 76, 2 mice). **c-e**, Same as Fig. 2**j** but for V2M S_Inc_ neurons (**c**), M2 S_Inc_ neurons (**d**), and M2 S_Dec_ neurons (**e**). **f-g**, Same as Fig. 2**l,n** but for M2 M_Inc_ (top) and M_Dec_ neurons (bottom).

**Supplementary Table 1.** The number of mice and neurons used in this paper.

| Experiment Types | Number of Mice | Number of Recording Sessions | Number of Neurons | Response Type |  |  |  | Figures |
| --- | --- | --- | --- | --- | --- | --- | --- | --- |
|  |  |  |  | Sensory | Motor | Dual | None |  |
| Retrograde tracing of ACC | 5 | - | - | - | - | - | - | Fig. 3a-d |
| Anterograde trans-synaptic (V2M) and retrograde tracing (DMS) | 1 | - | - | - | - | - | - | Fig. 5a,b |
| Effect of muscimol on neural activity | 1 | 1 | 33 | - | - | - | - | Fig. 1g |
| ACC inactivation (visual detection task) | 14 | - | - | - | - | - | - | Fig. 1h-k |
| V1 inactivation (visual detection task) | 10 | - | - | - | - | - | - | Fig. 1l-o |
| V2M inactivation (visual detection task) | 5 | - | - | - | - | - | - | Sup. Fig. 1a<sub>1</sub>-e<sub>1</sub> |
| M2 inactivation (visual detection task) | 4 | - | - | - | - | - | - | Sup. Fig. 1a<sub>2</sub>-e<sub>2</sub> |
| S1 inactivation (visual detection task) | 11 | - | - | - | - | - | - | Sup. Fig. 1a<sub>3</sub>-e<sub>3</sub> |
| ACC inactivation (stop signal task) | 7 | - | - | - | - | - | - | Fig. 7a-e |
| Photostimulation of ACC←V2M neurons (visual detection task, ChR2 group) | 3(9)* | - | - | - | - | - | - | Fig. 4a-e |
| Photostimulation of V2M axons in ACC (visual detection task, ChR2 group) | 3(12)* | - | - | - | - | - | - | Fig. 4a,c-e |
| Photostimulation of ACC←V2M neurons (visual detection task, control group) | 3(8)* | - | - | - | - | - | - | Fig. 5c,d |
| Pupil size and orofacial movement imaging (visual detection task) | 6 | - | - | - | - | - | - | Sup. Fig. 4 |
| ACC activity from untrained mice | 3 | 9 | 111 | 6 | 29 | 3 | 73 | Sup. Fig. 2 |
| ACC activity (visual detection task) | 21 | 26 | 550 | 96 | 102 | 27 | 325 | Fig. 1p-s, Fig. 2, Fig. 6a-c,g-o<br>Sup. Fig. 2, Sup. Fig. 5a,c,d<br>Sup. Fig. 6a-g |
| V1 activity (visual detection task) | 11 | 14 | 175 | 67 | 22 | 36 | 50 | Fig. 1p,q,t,u, Sup. Fig. 3<br>Sup. Fig. 4a,e-g<br>Sup. Fig. 5a,h-m |
| DMS activity (visual detection task) | 8 | 22 | 241 | 30 | 57 | 29 | 125 | Fig. 5c-e |
| V2M activity (visual detection task) | 2 | 4 | 76 | 13 | 7 | 2 | 54 | Sup. Fig. 7a-c |
| M2 activity (visual detection task) | 4 | 8 | 147 | 26 | 36 | 57 | 28 | Sup. Fig. 7a,b,d-g |
| Opto-tagging recording of ACC←V2M neurons (visual detection task) | 8 | 14 | 345 | 91 | 50 | 37 | 167 | Fig. 3e-h, Fig. 6d-f |
| Photostimulation of ACC←V2M neurons (visual detection task, ChR2 group) | 4 | 7 | 134 | - | - | - | - | Fig. 4f-i |
| Photostimulation of V2M axons in ACC (visual detection task, ChR2 group) | 2 | 2 | 96 | - | - | - | - | Fig. 4f,g,j |
| Photostimulation of ACC←V2M neurons (visual detection task, control group) | 2 | 4 | 72 | - | - | - | - | Fig. 4f,g |
| DMS activity recording with photostimulation of V2M axons in ACC (visual detection task) | 7 | 17 | 222 | - | - | - | - | Fig. 5f-h |
| ACC activity (stop signal task) | 5 | 9 | 162 | 23 | 57 | 27 | 55 | Fig. 7f,g |

\* Numbers in brackets indicate the number of sessions for optogenetics

**Supplementary Table 2.** Abbreviations of anatomical structures. All quantification of brain regions and boundaries is based on the Allen Reference Atlas (ARA).

| Brain structure | Abbreviation |
| --- | --- |
| Primary visual cortex | V1 |
| Anteromedial visual cortex | AM |
| Posteromedial visual cortex | PM |
| Anterolateral visual cortex | AL |
| Lateral visual cortex | L |
| Laterointermediate visual cortex | LI |
| Posterolateral visual cortex | PL |

**Supplementary Video 1. Orofacial movements of task-performing mouse**

IR imaging of four orofacial movements (pupil size, whisker movement, nose movement, and licking) from a mouse performing the visual detection task. Colored boxes indicate ROIs measured for each orofacial movement (black for pupil size, magenta for whisking movement, red for nose movement, cyan for licking). White ellipse, pupil boundary; white cross, center of pupil; green vertical lines, onsets of visual stimuli; blue dots, licking.

## Notes

### Competing Interest Statement

The authors have declared no competing interest.

